# Prediction of parsimonious and temporally sensitive sets of cell fate engineering transcription factors with IMCell

**DOI:** 10.64898/2026.06.10.731467

**Authors:** Emily Y. Su, Brandon Ly, Patrick Cahan

## Abstract

Transcription factor (TF) cocktails used in cell identity reprogramming protocols have largely been developed from experimental approaches. A handful of computational approaches have been reported, though have not been widely adopted by the scientific community. To standardize their use and assess their performance, we built CompForce, a platform that integrates these tools. Using CompForce, we found that existing computational methods offer modest improvements over differential expression on both synthetic and literature-curated data, and that their lackluster and inconsistent performance could be attributed to a reliance on local centrality metrics. To improve upon these methods, we developed IMCell, a prediction method that is inspired by the influence maximization problem. Unlike existing tools, IMCell returns optimized TF sets rather than ranked TF lists. We demonstrate that IMCell vastly out-performs existing tools, and further extend it to dynamic, stepwise contexts. The tools presented here are available in the R packages CompForce and IMCell.

## Introduction

Accurately and efficiently directing cell fate has long been an aspiration in the cellular engineering field that has been met with moderate success. At present, the differentiation, transdifferentiation, and reprogramming of numerous cell types have been reported. Most famously, the reprogramming of mouse embryonic and adult fibroblasts to pluripotent stem cells^1^ revealed the inherent plasticity of cellular identity and facilitated numerous studies that aimed to alter or guide cell fate^2–5^. Several strategies have been employed, including the use of small molecule mediators or growth factors and the over-expression or silencing of specific transcription factors (TFs). Regardless of method, the aim is to force the starting cell’s underlying regulatory circuitry to adopt a new configuration amenable for establishing and maintaining the target cell type identity.

To date, many reported cell fate engineering protocols have been developed experimentally. For example, overexpression-based protocols are typically discovered via systematic assessment of TF combinations, akin to the pipeline demonstrated by Takahashi and Yamanaka. Briefly, a candidate pool of TFs is narrowed by methodically testing the ability of sets of TFs to induce expression of target genes as read out by a reporter system. Unfortunately, such an approach is limited and unscalable for several reasons. First, the necessity of an initial TF candidacy list limits target cell types to those that are well-studied, such that key regulators of their identity are already known. Second, reporter systems and accurate experimental validation of identity may not be feasible for all cell types. Third, with roughly 2000 TFs and hundreds of cell types, the number of possible TF combinations is astronomical, making it infeasible to approach all such problems in a guess-and-check manner.

For those reasons, several computational prediction methods have been developed. A subset of these use transcriptomic data to rank TFs by the likelihood that their over-expression would successfully induce target cell type identity. The method proposed by D’Alessio et al. employs the Jensen-Shannon divergence (JSD) to identify TFs whose expression is central to a specific cell’s identity^6^. CellNet, in contrast, employs a network-based approach, in which a target’s cell type-specific gene regulatory network (GRN) is extracted from a background^7^. TFs are subsequently ranked based on a “network influence score” that accounts for expression of a TF and its targets in the desired cell type, the extent to which a TF and its targets expression are dysregulated, and the number of targets a TF has. Mogrify, another network-based approach, relies on related strategies, including ontogeny tree-based background selection, differential expression, and TF sphere of influence scoring^8^. However, despite the availability of these tools spanning several years, they have not been widely utilized to uncover novel or improve existing cell fate engineering protocols.

Similar to experimental approaches, several limitations likely prevent the widescale adoption of these computational tools. First, aside from smaller-scale benchmarking in their respective papers, there is limited literature on how well methods perform in comparison to each other and against less complex approaches such as differential expression of TFs. Existing benchmarking studies of computational tools unfortunately do not include nor focus on tools created specifically for the purposes of predicting TFs to target for reprogramming^9^. Second, not all methods are amenable to user flexibility, may be non-intuitive to implement, or may lack publicly available source code. For example, methods such as Mogrify do not allow for users to provide their own transcriptomic data and network information, and instead rely on FANTOM5, STRING, and MARA databases, limiting application. Third, existing methods return ranked lists of TFs, which mimic the candidate TF lists used in experimental approaches. It is typically unclear the number of top ranked TFs required to successfully induce target cell type fate, and further, it is unclear if the top ranked TFs work synergistically, have redundancy, or even work antagonistically. In other words, existing computational tools do not extract the minimal necessary and sufficient TF sets which experimental approaches attempt to uncover.

To address the first two limitations of computational approaches, we developed CompForce (“COMPutational tools FOR Cell fate Engineering”). CompForce is a tool in R that integrates multiple expression-based prediction approaches into a single platform, returning prediction results from individual methods as well as results from aggregation across methods. CompForce adds flexibility to each method, which includes allowing users to input their own expression datasets and networks, while also standardizing the input and outputs of each method, which allows for easy implementation and method comparison. Additionally, CompForce is built modularly, such that novel methods can be incorporated easily.

We used CompForce to benchmark existing methods and their aggregation against differential expression on two distinct classes of gold standards. First, we created a series of synthetic datasets via Dyngen^10^, and used CompForce to predict TF rankings across multiple transitions representative of directed differentiation, transdifferentiation, and reprogramming. We simulated perturbation of TFs based on these predictions and compared the resulting identities of simulated cells against the synthetic “wild type” target. Second, we compiled a series of literature-based gold standards, comprised of known TF sets capable of inducing specific transitions as well as TFs that have been shown to be unnecessary for the same transitions. Using five curated transcriptomic datasets, we applied CompForce and compared prediction results against these gold standards. To our surprise, we found that more complex methods that employ network structural information did not necessarily out-perform a pure differential expression approach.

Because existing network-based approaches often rely on centrality metrics to rank the importance of TFs in inducing specific cell fates, we further explored the predictive power of various centrality metrics and their robustness to noise. Using three separate literature-curated GRNs, we found that no single centrality metric we tested could distinguish literature-reported driver TFs (those capable of inducing a target cell fate) from non-driver TFs. To determine if a combination of centrality metrics or if the addition of expression-based metrics could distinguish driver and non-driver TFs, we trained separate Random Forest classifiers to make this distinction based on centrality metrics alone, expression-based metrics alone, and combined. We ultimately found that combinations of centrality metrics added little value to prediction, and that in fact expression-based metrics tended to have more predictive power.

Thus, we postulated that for networks to provide more utility to prediction, it was necessary to go beyond centrality metrics and consider more nuanced methods of measuring TF influence. We also wanted to address the third limitation of existing methods, and move beyond returning ranked lists to returning minimal TF sets. Taking inspiration from the network science field, we employed the Influence Maximization (IM) problem to our GRNs. Briefly, IM refers to the problem of finding a small set of nodes in a network that maximizes the spread of information^11^. It is often applied in the context of social networks, for example to find a subset of individuals to target such that an advertisement or product is spread to the largest number of people. In our context, we formulated the TF prediction problem to identify the minimal set of TFs needed to activate and repress expression of a collection of specific target genes. We present multiple variations on this method, which we have named IMCell, and demonstrate their superiority in identifying driver TFs using the CompForce platform. Finally, we demonstrate how IMCell can be easily applied to dynamic GRNs. This is particularly useful to identify sets of TFs that could be targeted across multiple steps of a differentiation protocol. IMCell is available as a package in R.

## Results

### CompForce is a platform for computational transcriptomic prediction tools

A handful of transcriptomic-based computational prediction tools have previously been developed. However, they vary in flexibility and required input, making fair comparison and benchmarking difficult. Additionally, from a user’s perspective, the lack of flexibility may limit application and ease-of-use. For these reasons, we developed CompForce, a platform that aims to standardize existing approaches and allow users to provide their own transcriptomic data (**Fig 1a**).

**Figure 1.**
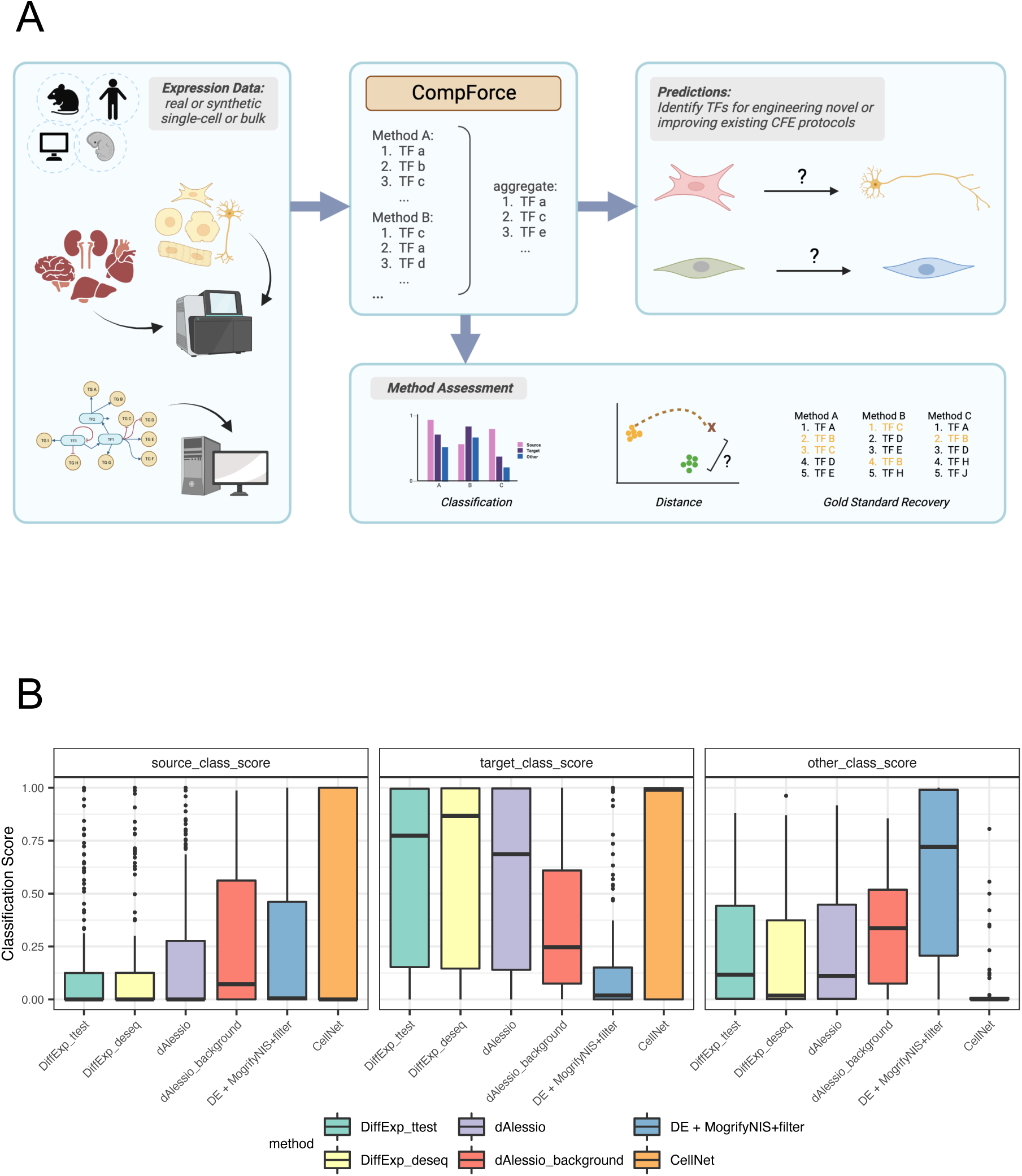
CompForce Platform and Synthetic Benchmarking. (A) The CompForce platform. CompForce takes as input transcriptomic expression data and optional network information. It then runs a series of computational prediction methods and aggregates results. The resulting predictions can be used to develop or refine cell fate engineering protocols, or be used for benchmarking purposes to assess prediction methods. (B) Classification scores representing how close simulations resembled source cell type, target cell type, and all other cell types. The plot shows simulation results from the overexpression of the top 8 TFs predicted by each method. 100 simulations were run for each prediction, and their resulting average is plotted.

CompForce takes as input single-cell or bulk transcriptomic data. At a minimum this data should be representative of starting and target cell states, but will often include background data. Users will also specify the starting and target cell states. Optionally, CompForce will take in a network structure, and users can further specify parameters specific to each approach.

From here, CompForce will run all or a user-specified subset of methods. First, as a baseline two forms of differential expression analysis are included: DESeq2^12^ and t-test. For these two approaches, TF expression in starting and target cell states are directly compared. DESeq2 is a differential expression analysis method for count data based on the negative binomial distribution. For the t-test approach, CompForce will first normalize and log-transform the data prior to the t-test. Both approaches then rank the importance of TFs based on adjusted p-value.

CompForce implements a version of the approach from D’Alessio et al. in which TFs are scored based on the JSD. In this approach, two pattern vectors are created. The first is the idealized pattern, in which elements are assigned a value of 1 at positions representing the target cell type and 0 at all other positions. The second is an observed pattern, which corresponds to the expression of a given TF across the dataset. CompForce includes two variations on this approach: in the first the data is limited to starting and target cell states, and in the second all non-target cell states are included and considered background. For both variations, TFs are then ranked based on the JSD between idealized and observed pattern vectors.

CellNet is a network-based approach in which TFs are ranked based on a “Network Influence Score” (NIS). In brief, this score is based on the number of target genes a TF has, the normalized and log-transformed expression level of the TF in the target state, and the extent to which the TF and its targets are dysregulated between starting and target states. The network CompForce employs in this approach is either supplied by the user or will be reconstructed using Context-Likelihood of Relatedness (CLR)^13^. In either case, cell-type specific networks are extracted prior to NIS computation and subsequent TF ranking.

Mogrify is a second network-based approach that similarly relies on inferring the importance of a given TF in the context of its location in a network. Mogrify’s approach relies on a number of strategies including background selection based on an ontogeny tree, differential expression via DESeq, and a network influence metric dependent on differential expression, out degree, and path length. Importantly, expression and ontogeny data are implemented from the FANTOM5 Consortium, and network influence is computed from the MARA and STRING networks (which results in two distinct network scores prior to Mogrify’s final ranking). Therefore, CompForce’s implementation of the Mogrify approach is not a direct application of the method. For example, though DESeq and final TF filtering are implemented exactly as is the original method, CompForce will instead use user-supplied expression data. Additionally, rather than two distinct network scores, only one is computed and is based on a network that is user-supplied, CLR-reconstructed, or from STRING. In this way, CompForce implements the metrics and approaches used within Mogrify without requiring the data from which it is originally built.

Finally, after all (or a user-specified subset) methods are run, CompForce can aggregate the results. Importantly, all of these methods return ranked lists of TFs based on some metric of importance. CompForce will use Robust Rank Aggregation (RRA)^14^ to combine these lists into an aggregated ranking. CompForce’s final output is thus individual ranked TF lists from each method and one aggregated ranked list.

### Benchmarking existing tools on synthetically generated data

To understand how well existing computational prediction tools perform, we applied CompForce to synthetically generated data. We used the Dyngen package to generate fifteen synthetic network models (**Supplementary Table 1**). Of these, five were bifurcating trajectories, five were trifurcating trajectories, and five were consecutive bifurcating trajectories. The network models spanned varying numbers of TFs (20-200), target genes (50-1100), and housekeeping genes (50-200). Each simulated and extracted expression dataset contained 1000 cells in total. Bifurcating trajectories include one progenitor state and two terminal states, while trifurcating and consecutive bifurcating trajectories include one progenitor state and three terminal states. This means that each bifurcating model covers six transitions (differentiation, transdifferentiation, and reprogramming) and each of the trifurcating and consecutive bifurcating models covers twelve transitions. In total, this equates to 150 total transitions to assess.

For each transition, we ran CompForce to generate ranked TF lists for each method. To assess these predictions, we modified the Dyngen approach to simulate the overexpression of the top ‘X’ ranked TFs generated by each method (X = 2, 3, 5, 8, 10). Each simulation was run 100 times, and the results of the 100 simulations were compared against the wild-type target using SingleCellNet^15^ (SCN) (**Supplementary Fig. 1a**). Interestingly, we found that network usage did not guarantee more accurate TF predictions. For example, CompForce’s implementation of Mogrify consistently performed worse than differential expression, with overexpression simulations resulting in lower target classification scores and higher off-target classification scores. Additionally, though CellNet performed the best amongst all methods, its performance was extremely variable, demonstrating a high degree of inconsistency with its predictions. Notably, performance for most methods began peaking at X=8 (**Fig. 1b**), which we later found was more TFs than necessary to drive the synthetic cell identity transitions. At this and lower values of X we found that overall, computational prediction methods demonstrated marginal improvement over differential expression (**Supplementary Fig. 1b**), and that predicted TFs returned by the different methods varied widely (**Supplementary Fig. 2**).

### Benchmarking existing tools on literature-curated data

To generate a more comprehensive picture on method performance, we next turned our attention to benchmarking on real world data. To this end we curated two single-cell expression datasets (a broad dataset encompassing 26 cell types and a neural-focused dataset) and three bulk expression datasets (a broad dataset encompassing 14 cell types, a blood dataset, and a neural dataset). For this comparison we also ensured that all prediction methods utilized the same network for a given dataset. In this way, we were able to isolate and test each method’s approach fairly. Networks for single-cell datasets were reconstructed using CLR. Networks for bulk datasets were curated from Enrichr^16,17^.

We next compiled positive and negative gold standards based on TF cocktails known to induce specific cell fate transitions (**Supplementary Table 2**). Positive gold standards were comprised of TFs in reported cocktails. Negative gold standards were comprised of TFs that had previously been tested but were demonstrated as unnecessary and that also did not appear in the corresponding positive gold standard. In total, we collected positive and negative gold standards for 19 cell identity transitions.

We then used CompForce to make TF predictions via each method, and assessed performance by counting the number of positive and negative gold standard TFs recovered (**Fig. 2, Supplementary Fig. 3**). Similar to our results with synthetic data, there were clear examples where using network information didn’t necessarily improve performance when compared to differential expression. For example, our implementation of Mogrify tended to recover fewer positive gold standard TFs. Further, while CellNet performed relatively well amongst the prediction methods, its improvement in recovering positive gold standard TFs over non-network based methods (including differential expression) was marginal and (similar to other methods) predicted a significant portion of negative gold standard TFs.

**Figure 2.**
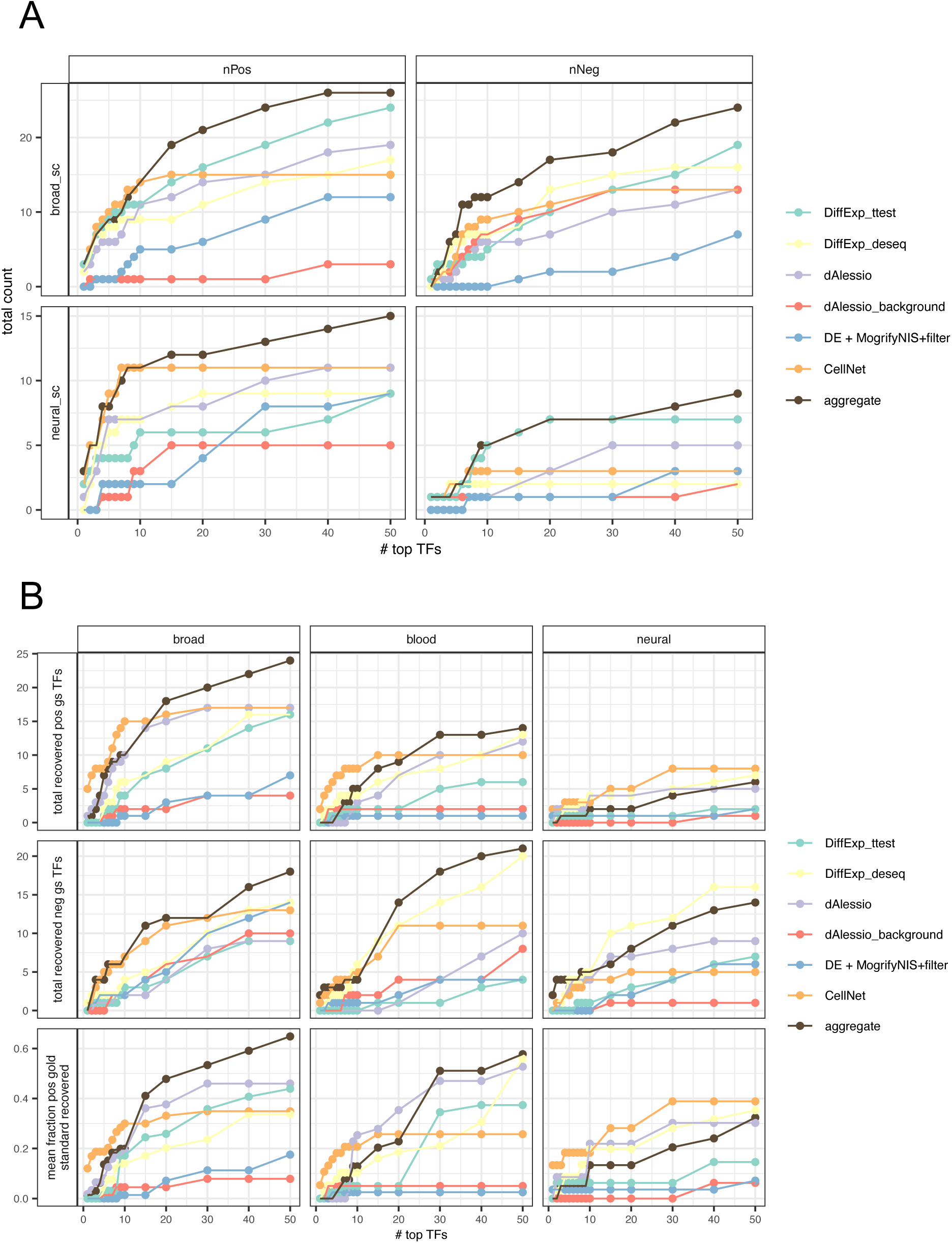
CompForce Literature-Curated Benchmarking. (A) The number of positive and negative gold standard TFs recovered by each method across the two single-cell datasets. (B) The top two rows show the number of positive and negative gold standard TFs recovered by each method across the three bulk datasets. The bottom row indicates the fraction of the positive gold standard TFs recovered.

Importantly, we also assessed the extent to which positive gold standards could be predicted by each method. Alarmingly, we found that all methods struggled in recovering full gold standard cocktails even when looking at the top 20 or 30 ranked TFs (**Fig 2b. Supplementary Fig. 4**). Ultimately, this suggests that existing computational methods provide limited insight in identifying TFs important in driving specific cell transitions, and that significant improvements must be made for such tools to be broadly useful in aiding the development or refinement of cell fate engineering protocols.

### Common centrality metrics are not robust and fail to identify driver TFs

We next sought to understand why computational prediction methods that incorporate network information did so with marginal performance benefits. As described above, network-based methods incorporate expression information with specific centrality metrics to rank TFs. Because the networks utilized in these prediction tools are likely incomplete and not fully accurate, we first began by testing the robustness of various common centrality metrics to noise in real biological networks. To do this, we curated interactions from the mouse and human networks from DoRothEA^18^, TRRUST^19^, and RegNetwork^20^. To these six networks, we progressively added 10%, 20%, 50% or 75% noise by randomly adding in or removing edges, and generated 50 perturbed networks for each noise level and network. This process was repeated with the synthetic network models. We then compared how TFs ranked in centrality between the perturbed and original networks (**Supplementary Fig. 5-6**). We found that all of the centrality metrics we assessed were affected by noise and that there was no stand out robust centrality metric across all networks.

More importantly, we then looked at the ability of common centrality metrics in distinguishing driver and non-driver TFs. In our case, we are defining driver TFs as those that that appear in our curated literature-based positive gold-standard, or in other words TFs that have been reported in successful TF reprogramming cocktails. For this analysis we looked at mouse networks from DoRothEA, TRRUST, RegNetwork, and Enrichr (which we refer to as ChIP-X). We then compared the centrality ranks of positive gold standard TFs (i.e. driver TFs), negative gold standard TFs, and all other TFs (**Supplementary Fig. 7**). Unsurprisingly, we found that no single centrality metric was able to distinguish driver from non-driver TFs.

We then asked if a combination of centrality metrics could make this distinction. To this end, we employed a classification approach. We began by reconstructing starting and target cell type networks across the gold standard transitions. We then assembled a feature matrix in which features were centrality metric (and ranks) in either the source or target network and observations were driver and non-driver TFs (including both negative gold standard TFs and randomly selected “other” TFs) (**Supplementary Fig. 8a**). We then trained ten Random Forest (RF) classifiers on ten sampled training sets, holding out 20% of the data each time for validation. In addition to validating on the held-out data, we also tested on randomly selected “other TFs” which were labeled as negative, or non-driver TFs. Based on precision, recall, and balanced accuracy we found that a combination of centrality metrics from both source and target cell type networks was still not able to clearly distinguish driver from non-driver TFs (**Fig. 3a-b**).

**Figure 3.**
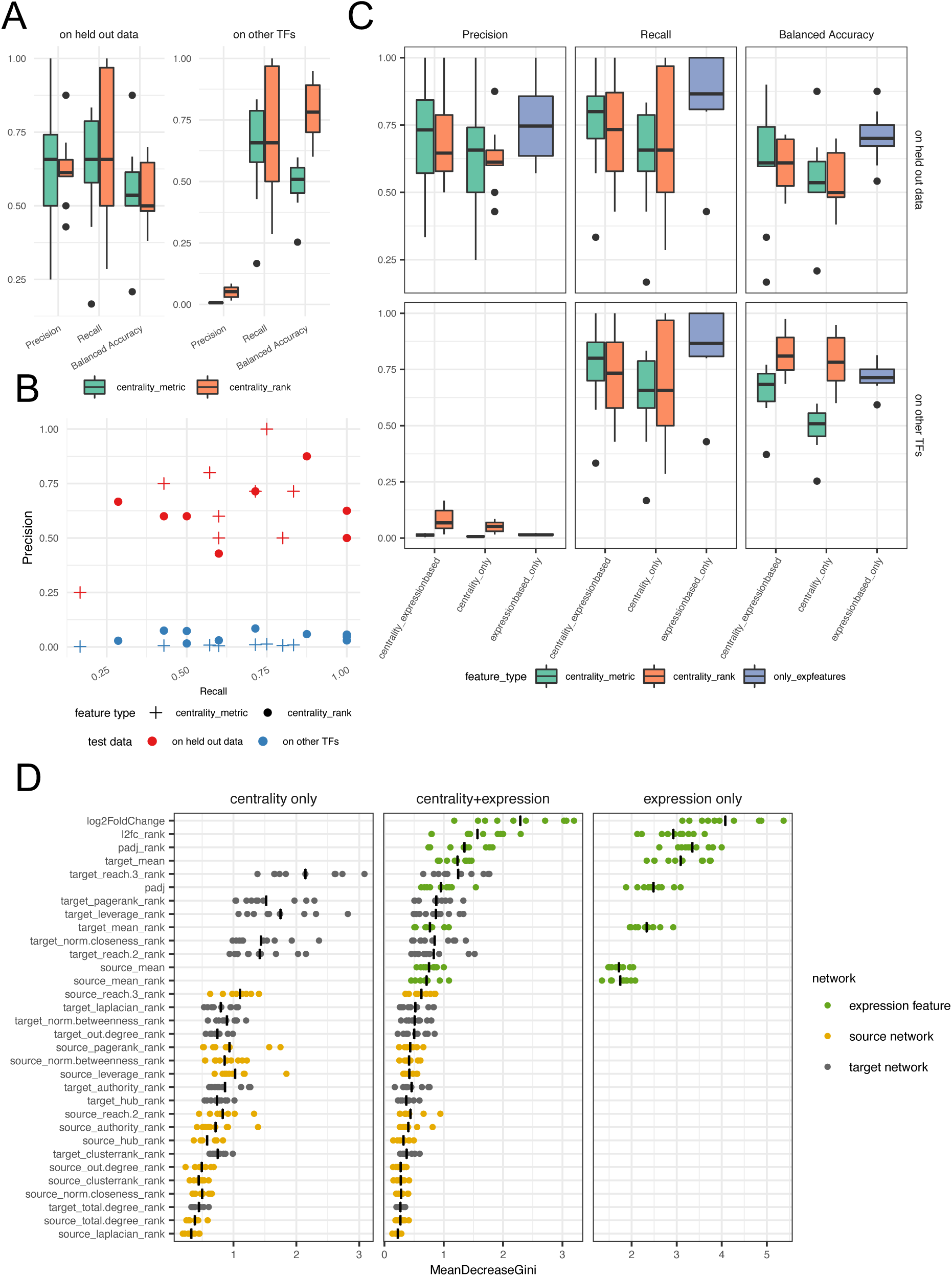
Predictive Power of Centrality Metrics. (A) Precision, recall, and balanced accuracy of RF classifiers trained on centrality metrics and centrality ranks. The left plot corresponds to validation on held-out gold standard data. The right plot corresponds to validation on the combination of held-out gold standard data and randomly selected other TFs (which are given a negative label). (B) Precision and recall of the RF classifiers trained on centrality metric and centrality rank. (C) Precision, recall, and balanced accuracy of RF classifiers trained on centrality features only, expression-based features only, and combined features. (D) Importance of each feature in the RF classifiers. Mean is indicated by the vertical bar, and features are ordered by importance.

Finally, we assessed the extent in which using the same centrality metrics improved classification over a purely expression-based feature matrix. To this end, we built an additional two types of feature matrices. The first included only expression-based features: mean expression in source and target cell type, mean expression rank in source and target cell type, adjusted p-value of the differential expression (DE) based on DESeq, rank of the differential expression adjusted p-value, the log-fold change in expression, and the rank of the log-fold change in expression. The second included both the centrality and expression-based features. Based on precision, recall, and balanced accuracy, we once again found that the combined centrality and expression-based classifiers did not out-perform the purely expression-based classifiers (**Fig. 3c, Supplementary Fig. 8b**).

Further, we ranked the importance of each feature in each classifier by decreasing Gini index and found that in general expression-based features such as log-fold change and adjusted p-value tended to be more important than centrality metrics (**Fig. 3d**). Interestingly, out-degree importance was very mid-tier, suggesting that a TF’s number of targets is not indicative of its ability to induce a particular cell fate transition, and further suggesting that its inclusion in computational prediction methods is likely unnecessary. The centrality metrics that ranked higher in importance tended to measure global influence as opposed to local influence. For example, reach-3 ranked highest and refers to the number of unique targets in a given TF’s 3-neighborhood.

### IMCell employs influence maximization to predict TF sets

We next aimed to improve upon existing computational prediction tools in two ways: first, by moving away from a reliance on local centrality metrics, and second, by identifying TF sets rather than ranked lists. As described in the previous section, local centrality metrics do not offer much additional predictive power in comparison to differential expression analysis. Further, existing methods return ranked lists rather than sets, and it is typically unclear how many top ranked TFs are necessary to dysregulate for a given cell identity transition. The relationship between these top ranked TFs is also unclear. In the ideal scenario, top ranked TFs should be synergistic; in the unideal scenario they may be redundant or even antagonistic. Of the existing approaches, only Mogrify broadens the target gene space by filtering out TFs that share 98% of their targets. However, this strategy is still limited given the high threshold and the fact that TF influence is evaluated at the individual TF level rather than as TF sets, which obscures synergistic, redundant, and antagonistic effects amongst TFs.

To address these limitations, we developed IMCell (**Fig. 4**), a computational prediction tool that takes inspiration from the influence maximization (IM) problem^11^. Briefly, in the IM framework, a node that is activated in a network adopts a particular behavior. Once a node is activated, it then has the opportunity to activate its neighbors with a given probability. The IM problem focuses on extracting the smallest seed set of *k* nodes whose activation maximizes the number of nodes activated in the network. In other words, the focus of the IM problem lies in maximizing the “spread” or “influence” of a set of TFs over a network. Since its conception, numerous variations on the IM problem have been developed, including for example, polarity-related influence maximization (PRIM)^21^. Notably, PRIM extends the problem to signed networks, such that nodes can have either positive or negative relationships between each other.

**Figure 4.**
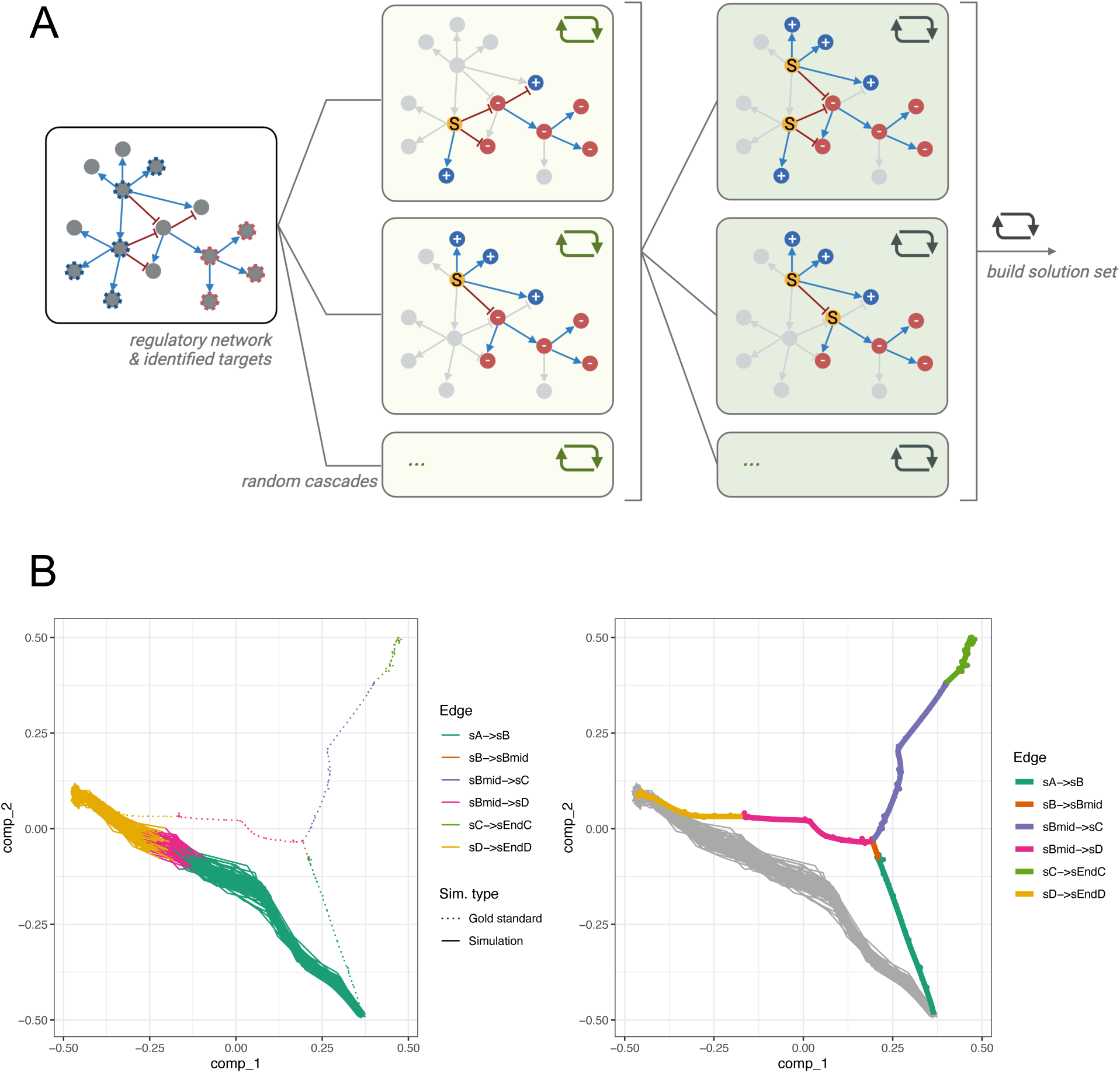
IMCell. (A) IMCell aims to maximize the activation and repression of a target set of genes. To do this, it employs a greedy Monte-Carlo approach in which iterations of random cascades are run to build the optimal solution seed set. (B) An example of 100 simulations in which an IMCell-predicted TF set is overexpressed. In the example, a TF set is predicted for the state C to state D direct conversion. The wild type trajectory and populations are shown as a dotted skeleton (left) and bold colored (right).

In IMCell, we extend the PRIM framework in three ways. First, while IM and PRIM both aim to maximize spread across all nodes in the network, our approach maximizes spread across specific target gene sets corresponding to a given genetic program. Second, PRIM focuses solving the positive or negative IM problems (PIM and NIM, respectively) in which the goal is to maximize positive or negative influence. IMCell, however, aims to maximize both, striving to activate a target program while repressing other off-target programs. Third, while the IM and PRIM problems view all nodes with equal weight, IMCell allows for optional node weighting based on transcriptomic expression data when computing spread. Based on this framework, IMCell then employs a greedy, Monte-Carlo based approach to predict a TF set capable of inducing specific cell fate transitions.

### IMCell out-performs existing computational prediction tools

We assessed the performance of IMCell utilizing CompForce and the same synthetic datasets described previously. We tested four variations of IMCell: (1) IMCell_CELF: a version that mimics the original IM problem, limiting the network to activating edges only and solved using the cost-effective lazy forward algorithm (CELF); (2): IMCell: the base version of IMCell; (3): IMCell_repressorwins: a version in which repressive interactions take precedence over activating interactions; (4): IMCell_expweighted: a version in which nodes (and spread) are weighted by expression data.

To assess IMCell’s performance, we benchmarked against existing tools using the CompForce platform and the datasets described previously. Importantly, because IMCell returns a TF set while other methods return ranked TF lists, we compared IMCell predicted sets to the top X TFs returned by each method, where X represents the number of TFs in the IMCell set. For example, if IMCell returned a set containing three TFs, then we would compare the result against the top three TFs returned by all other methods. Importantly, we found that for these datasets, IMCell returned 1-3 TFs per set. For the synthetic datasets we assessed performance by simulating the overexpression of predicted TF sets, using SCN and Euclidean distance to measure how closely simulated cells resembled the wild-type target population.

We found that overwhelmingly, IMCell and its variations out-performed all other prediction tools with simulated cells exhibiting high target classification score, low off-target classification score, and low distance to wild-type population (**Fig. 5a-b, Supplementary Fig. 9**). Amongst the four IMCell variations we tested, IMCell, IMCell_repressorwins, and IMCell_expweighted performed best, indicating the importance of considering repressive edges when measuring influence. Interestingly, we also found that all methods had a more difficult time predicting TFs for reprogramming transitions (i.e. terminal state to progenitor state). This was especially the case for CellNet. We speculate that this is largely due to the nature of the synthetic data, in which the network is designed modularly such that the progenitor state easily gives rise to downstream states. However, even without considering reprogramming transitions, IMCell and its variations drastically outperformed all other methods. Additionally, we found that even with varying levels of added noise to the input network, IMCell still outperformed CellNet and our implementation of Mogrify at equivalent noise levels, demonstrating its robustness to network inaccuracy (**Fig. 5c**).

**Figure 5.**
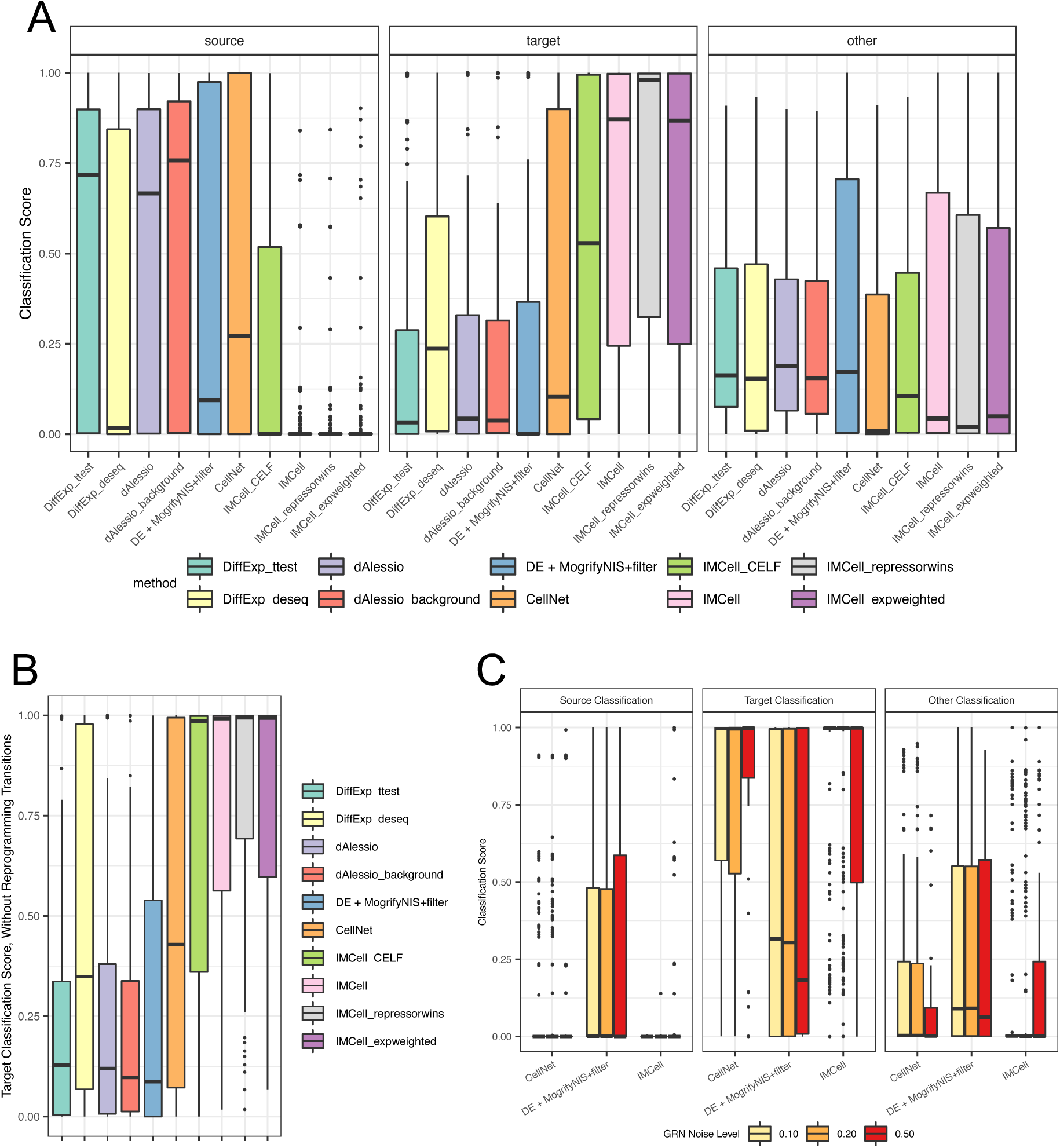
IMCell Benchmarking. (A) SCN classification scores detailing how closely simulated cells overexpression predicted TF sets resemble the source, target, and other populations. For non-IMCell methods, TF sets were defined as the top X TFs returned by each method, where X is the number of TFs in the IMCell result. (B) Target classification scores disregarding reprogramming transitions. (C) Classification scores corresponding to network-based methods at different levels of network noise.

### IMCell can be extended to predict stepwise cell fate engineering protocols

IMCell can be extended to a dynamic context in which the goal of engineering a particular cell fate transition is broken down in a stepwise manner (**Fig. 6a**). This is accomplished by first applying Epoch^22^, a single-cell transcriptomic GRN reconstruction tool, to the input network. Epoch has the capability to extract a dynamic, or temporal, network from a static network. From here, sequential rounds of IMCell are run using each epoch subnetwork as input. This results in series of predicted TF sets, which correspond to a stepwise progression of a cell fate engineering protocol.

**Figure 6.**
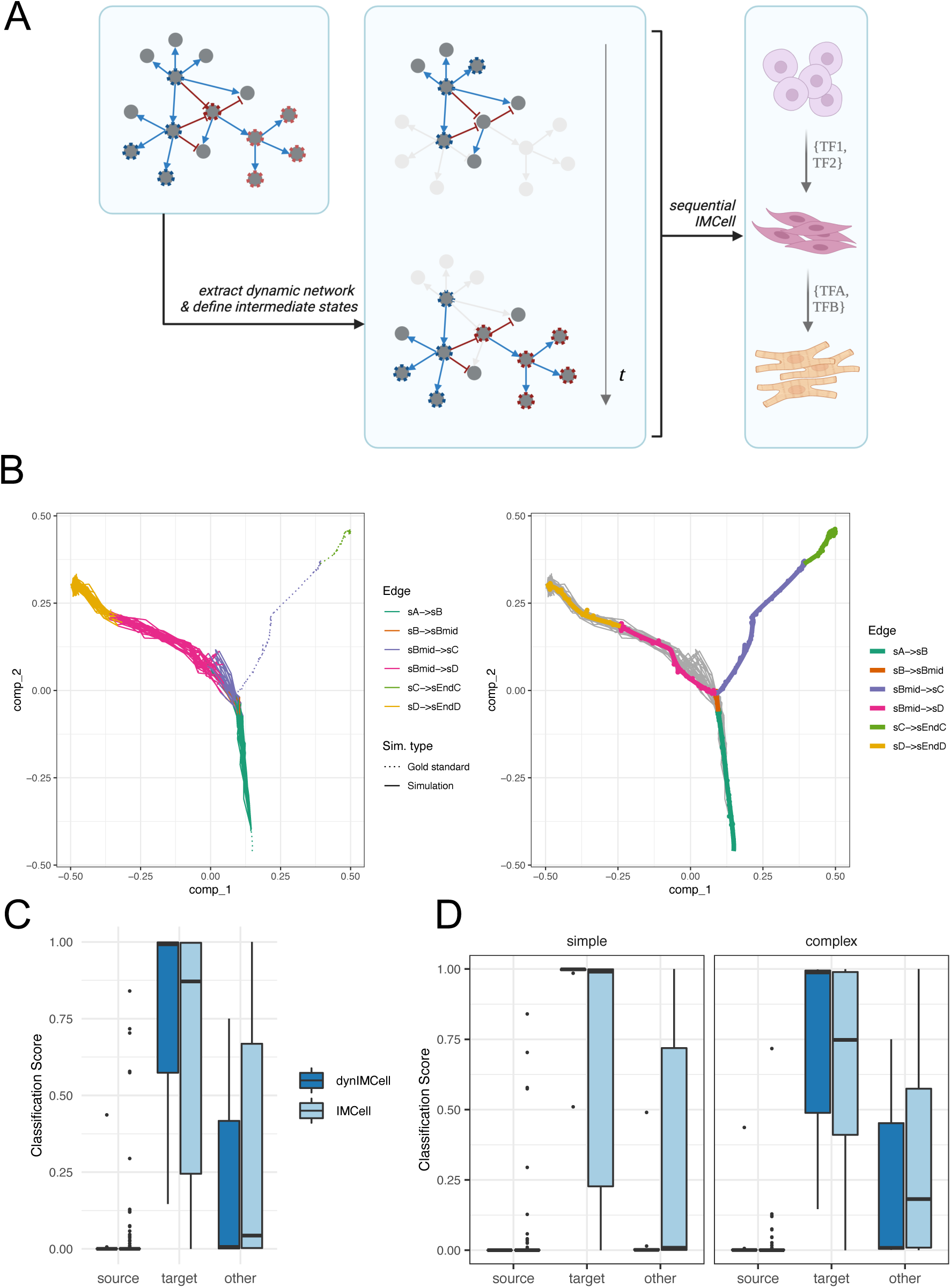
Applying IMCell to Dynamic, Stepwise Fate Engineering. (A) Schematic detailing how IMCell can be applied to stepwise fate engineering. IMCell employs Epoch to break the static network into a dynamic network. Sequential rounds of IMCell are run to create a stepwise protocol. (B) An example of 100 simulations in which a stepwise protocol predicted by IMCell was applied. Rather than a direct jump from source to target cell type, the cells are guided through an intermediate state, mimicking the wild-type trajectory. (C) Source, target, and other classification scores of the results of overexpression simulations in which protocols were predicted by IMCell versus the dynamic application of IMCell. (D) Breakdown of the classification results based on if the trajectory was a bifurcating trajectory (simple) or consecutive bifurcating trajectory (complex).

By applying IMCell in a stepwise context, we were able to guide differentiation through specific intermediate states (**Fig. 6b**). We then aimed to understand the utility of a stepwise differentiation strategy versus a “jumping” single-step transition. To do so, we focused on the differentiation transitions (i.e. from progenitor to terminal states) within the bifurcating and consecutive bifurcating synthetic models described previously. We used IMCell to predict TF sets for each transition and further applied the dynamic version of IMCell (“dynIMCell”) to predict TF sets that would guide differentiation through the intermediate branch points. We then simulated the overexpression of the predicted sets and assessed the resulting cells using SCN. We found that there was a modest benefit to the stepwise strategy, with improvements in target classification score (**Fig. 6c**). Interestingly, we found that this benefit was more pronounced for the more complex, consecutive bifurcating trajectories (**Fig. 6d**). These results suggest that more complex cell identity transitions or transitions between very distinct identities could ultimately benefit from a stepwise engineering approach guided by a tool like IMCell.

## Discussion

Computational prediction tools offer a solution for uncovering key TFs that could be targeted for the purposes of inducing cell identity transitions. Unfortunately, there are only a handful of such tools that exist. Further, existing tools are limited in their flexibility and predictive power, preventing their widespread application in the context of cellular fate engineering. Because of this, experimental approaches involving TF screens remain the standard in developing fate engineering protocols. And notably, as experimental methods advance, larger-scale screening approaches have proven to be incredibly fruitful, for example in programming human pluripotent stem cells toward diverse tissues^23^.

Computational prediction methods currently focus on ranking TFs by some measure of importance, which is dictated either by specific expression in the target cell type or influence in a regulatory network. Producing ranked lists of candidate TFs is unideal for two reasons. First, it is unclear how many of the top ranked TFs are necessary to induce a particular transition. Second, the relationship between top ranked TFs is typically unknown. Despite this, such approaches could be a complementary strategy that could be applied to simplify experimental screens. However, subpar performance ultimately limits their utility in this context.

With the aim of advancing computational approaches to identify key regulators of cell identity transitions, we built CompForce, a platform that integrates and standardizes existing prediction tools. CompForce allows users to input their own expression data and networks and will return predictions from both individual methods as well as an aggregated result. Importantly, CompForce is built modularly, such that additional methods can be easily incorporated over time.

We used CompForce to benchmark existing methods against differential expression on both synthetic and literature-curated data. Of note, though literature-curated data is ideal for its direct relevance, its use comes with several caveats. First, the networks utilized are likely inaccurate or incomplete and could skew the performance of network-based methods. Second, gold standards can be biased given that candidate TFs used in experimental approaches were likely chosen based on knowledge of their specific expression or regulatory role in the target cell type. For these reasons, what we deem as an inaccurately predicted TF may just be a TF that has not been previously tested or tested in the right combination of factors. Thus, our use of synthetic data can be considered a more direct approach of assessing TF predictions.

Ultimately, we found that even though computational prediction methods were developed for the purpose of identifying key regulators, their improvement over differential expression was modest at best and could be inconsistent across datasets and transitions. Interestingly, the inclusion of network information did not necessarily improve prediction. Our analyses suggest this is due to the limited predictive power of centrality metrics in ranking influential TFs in the context of cell fate reprogramming. Indeed, centrality metrics did not significantly improve the distinction between driver vs. non-driver TFs over purely expression-based metrics.

We then focused on improving upon these methods by developing IMCell, which does not rely on local centrality metrics and returns optimized TF sets rather than ranked lists. IMCell is inspired by the influence maximization problem often applied to social networks but extends the framework in several ways. First, IMCell aims to maximize spread across specific target gene sets corresponding to specific genetic programs. Second, IMCell considers both the activation and repression of target genes in determining influence. Third, IMCell allows for optional node weighting, refining the predicted spread and influence. We demonstrated that IMCell out-performs existing computational approaches, is robust to network inaccuracies, and further demonstrated that it can be utilized in a dynamic scenario to predict a series of TF sets for stepwise differentiation protocols.

Importantly, IMCell differs from existing methods in its required inputs and approach, and it does not strictly require transcriptomic data. While other methods search for larger-scale transcriptomic differences between starting and target cell types, IMCell’s premise is centered around the activation and repression of specified gene sets. For example, instead of identifying all differences between two cell types, which likely includes multiple genetic programs and could dilute the power of a given tool, IMCell can take in a single set of target genes (even without corresponding transcriptomic data) involved in a specific cellular function and search for the optimal TF set to activate these targets. Thus, this makes IMCell particularly useful in the targeted engineering of specific cell functions, which other tools would not be able to accomplish.

In addition, we envision that IMCell and its dynamic application will be useful in answering several fundamental questions. For example, such a tool can aid in exploring the extent to which cell identity transitions are plastic, allowing researchers to explore multiple routes (i.e. TF sets) of inducing the same transition. Further, the dynamic application of IMCell can aid researchers in assessing the benefits or drawbacks of direct reprogramming versus the stepwise “nudging” of cell identity. Our own initial analyses suggest that a stepwise approach may be advantageous in more complex transitions.

Finally, we note that the existing tools we assessed take a transcriptomic-based approach toward prediction, which may be limited by the nature of the data itself. For example, the integration of epigenetic information can likely illuminate roadblocks in reprogramming or highlight easier paths of transition. Additionally, it remains unclear if any computational approach could ever exceed a pure large-scale genetic screen. However, given the scale of the problem, computational approaches remain a promising complementary approach toward advancing cell fate engineering.

## Methods

### The CompForce platform

CompForce is a platform for computational fate determinant prediction tools that rank influential TFs in cell identity transitions. At the core of CompForce is the ‘CompForce’ class. This is an object that stores raw (counts) expression data, corresponding metadata, optional network information, individual method specific parameters (such as species, TFs, etc.), and results. Currently the CompForce platform supports the following methods: DESeq2, t-test, an implementation of the method reported by D’Alessio et al., an implementation of Mogrify, and CellNet. CompForce is structured modularly such that novel methods can be easily incorporated into the platform. It is available as a package in R.

### Synthetic dataset generation and simulation

We used the Dyngen package in R to generate fifteen synthetic network models. Five were bifurcating trajectories, five were trifurcating trajectories, and five were consecutive bifurcating trajectories. The network models spanned varying numbers of TFs (20-200), target genes (50-1100), and housekeeping genes (50-200). 100 simulations were run for each model, cell profiles were sampled, and random variation emulating a sequencing experiment was added. Each simulated and extracted expression dataset contained 1000 cells in total. Specific details of each network model can be found in the supplementary information.

To run overexpression simulations, we modified the Dyngen perturbation pipeline, altering the starting cell state, removing the burn-in time, adjusting the basal state of overexpressed TFs, and modifying propensity functions of overexpressed genes such that their transcription was continuous. All simulations were run 100 times and resulting cells were compared against wild-type using SCN and Euclidean distance.

### Real-world network curation and cell-type specific networks

Mouse and human networks were curated from DoRothEA and filtered for reliability. Interactions in DoRothEA are graded based on reliability. We limited our networks to interactions that were graded “A”, which indicate support from literature-curated resources, ChIP-seq peaks, inference from gene expression, and TF binding motifs on promoters. Mouse and human networks were curated from TRRUST as is. Mouse and human networks were curated from RegNetwork by filtering for TF-target gene interactions and omitting protein-protein interactions. Curated networks were made cell-type specific by limiting interactions to active genes in the corresponding datasets.

### Centrality and Random Forest analysis

For each curated network, 50 perturbed networks were generated at each noise level (10%, 20%, 50%, 75%) by randomly adding or removing edges. Centrality metrics were computed for each TF in each network, and centrality ranks for each TF were computed by ranking them by the corresponding metric. These results were compared against those from the unperturbed networks, and change was measured by percent change in rank, Jaccard similarity, and Kendall’s tau.

Random Forest classifiers were built from a feature matrix that detailed centrality metrics and ranks from CLR-reconstructed networks representative of gold standard transitions (from both starting and target cell types). Expression-based features were computed using DESeq2.

### IMCell

IMCell uses a greedy, Monte-Carlo (MC) approach to predict a TF set that maximizes influence or “spread” in a given network. At a minimum, IMCell requires a directed network structure as input (weighted or unweighted), though various other inputs and parameters can be specified by the user including expression and metadata, specified target genes of interest, limitations on resulting set size, limitations on TF search radius, limitations on marginal spread, amongst others.

At the core of IMCell is the random cascade. Given a directed finite network, a random cascade is initialized by activating a seed set of nodes. A node that is activated adopts a particular behavior (i.e. a gene can turn “on” or “off”). Additionally, once activated, the node now has the opportunity to activate its targets with a specific probability. In other words, the adopted behavior has the opportunity to be carried forward. Each edge has only one chance of firing. Should more than one edge incident to the same target node successfully fire, the winner is chosen by uniform random sampling. This activation spreads through the network to create a random cascade. The “spread” or influence is then calculated as the total number of targets that are correctly turned on or off in the network.

The rules of the random cascade (e.g. regulatory functions) vary depending on the variation of IMCell that is run. However, all variations utilize the same principle to build a predicted TF set. Namely, IMCell will start by employing a MC approach and simulate numerous random cascades to estimate each individual TF’s expected spread. IMCell will then add the TF with the largest expected spread to the solution set. In the next iteration, the random cascades are simulated again, but this time seeding with the union between the solution set and one additional TF. These iterations continue until one of three conditions is satisfied: the expected spread is maximized, the solution set size has reached a specified size limit, or the marginal spread (i.e. difference in expected spread between iterations) is sufficiently small. IMCell runs cascades in parallel so as to improve efficiency.

With respect to target genes of interest (whose activation or repression is used to compute spread), users have three options. They may opt to manually specify target genes, which is particularly useful for those looking to activate and repress a specific program or set of genes. Alternatively, users can utilize the built-in *find_differential_nodes* function (which relies on differential expression) to identify ideal subsets of genes for activation and repression between starting and target cell states. Finally, if left NULL, IMCell will compute spread across the entire network.

Users can also alter how edge firing probabilities are computed within the network. By default, IMCell will compute probabilities by normalizing by the in-degree at each node. Otherwise, IMCell can compute probabilities by normalizing based on total in-degree of the whole network or alternatively simply assign an equal probability across all edges. Further, users can alter an edge probability multiplier, should they find that the average path length is insufficient due to high edge density.

Users can also provide parameters that can refine the search. For example, the maximum number of TFs allowed in the solution set can be specified. Further, the search can be limited by a minimum marginal spread, in which IMCell will terminate early and return a truncated TF set if spread does not increase by the specified amount between iterations.

### TF scope in IMCell

The TF search radius can be limited in IMCell. This is particularly useful for large networks with a large number of TFs, on which predicting a TF set would be slow. Users can specify which TFs to limit the search to, or they can utilize IMCell’s built-in capabilities. IMCell implements a combination of two approaches to limit the TF scope. The first approach is via differential expression using DESeq2. IMCell will extract the top differentially expressed TFs based on adjusted p-value and log-fold change. The second approach is via template matching and pattern testing, with IMCell extracting the top TFs based on adjusted p-value and correlation. IMCell will take the union of the results from the two approaches to limit the TF scope.

### Common variations on IMCell

In this work, we assessed four variations of IMCell, though numerous other variations exist dependent on user-specified parameters. A summary of their differences is as follows. One of the variations, which we’ve labeled IMCell_CELF, mimics the original IM problem. In this variation, the input network is first limited to activating edges only. In other words, repressive interactions are ignored. From here, rather than running the previously described greedy approach, IMCell will implement the CELF algorithm.

A second variation, which we’ve labeled IMCell_repressorwins, follows the base version of IMCell with one difference: in given random cascade, should multiple edges fire successfully, repressive edges take precedence. In other words, if a repressive edge fires successfully, the target node will be repressed regardless of if activating edges also fired successfully.

The final variation, which we’ve labeled IMCell_expweighted, is an expression-weighted version of IMCell. This was created with the intention of utilizing expression data to provide supporting information for when the input network structure is suspect or when users want to prioritize transcriptomic information. In this variation, nodes are assigned weight based on their expression in starting and target cell states. Two weighting options can be implemented in IMCell. In the first option, weighting is proportional to mean expression in the target cell state. Otherwise, a differential weighting scheme is applied in which source cell state expression is z-scored in the context of the target cell state.

In our benchmarking analyses, we used *find_differential_nodes* to specify target nodes of interest. We further tested a minimal marginal spread of ten and twenty percent of the target node space, though there was no discernible difference in result. We computed edge weight probabilities by default in-degree, and did not apply any edge weight multiplier. TF scope was only limited for the 6 largest synthetic network models.

## Data and Code Availability

The CompForce and IMCell R packages are openly available on GitHub (https://github.com/CahanLab/compforce and https://github.com/CahanLab/IMCell). The exact software versions used in this study are archived on Zenodo: CompForce v0.1.0 (DOI: 10.5281/zenodo.20563029) and IMCell v0.1.0 (DOI: 10.5281/zenodo.20563020), each corresponding to release tag v0.1.0 of the respective repository. The transcriptomic datasets supporting this study—two single-cell and three bulk datasets, together with their cell-type-specific gene-regulatory networks, transcription-factor lists, and positive and negative gold-standard cocktails—and the synthetic Dyngen network models are archived on Zenodo (DOI: 10.5281/zenodo.20563179). Dataset provenance is summarized in Supplementary Tables 1 and 2.

## Author Contributions

E.Y.S. and P.C. conceived and designed the study. E.Y.S. developed the CompForce and IMCell software and performed the computational analyses. B.L. contributed to [specify contribution]. E.Y.S. and P.C. wrote the manuscript with input from all authors. P.C. supervised the study and acquired funding. [Authors: please confirm and finalize contributions, e.g., using CRediT taxonomy.]

