## Supplemental figures and tables for "Prediction of parsimonious and temporally sensitive sets of cell fate engineering transcription factors with IMCell"

### Slide 1
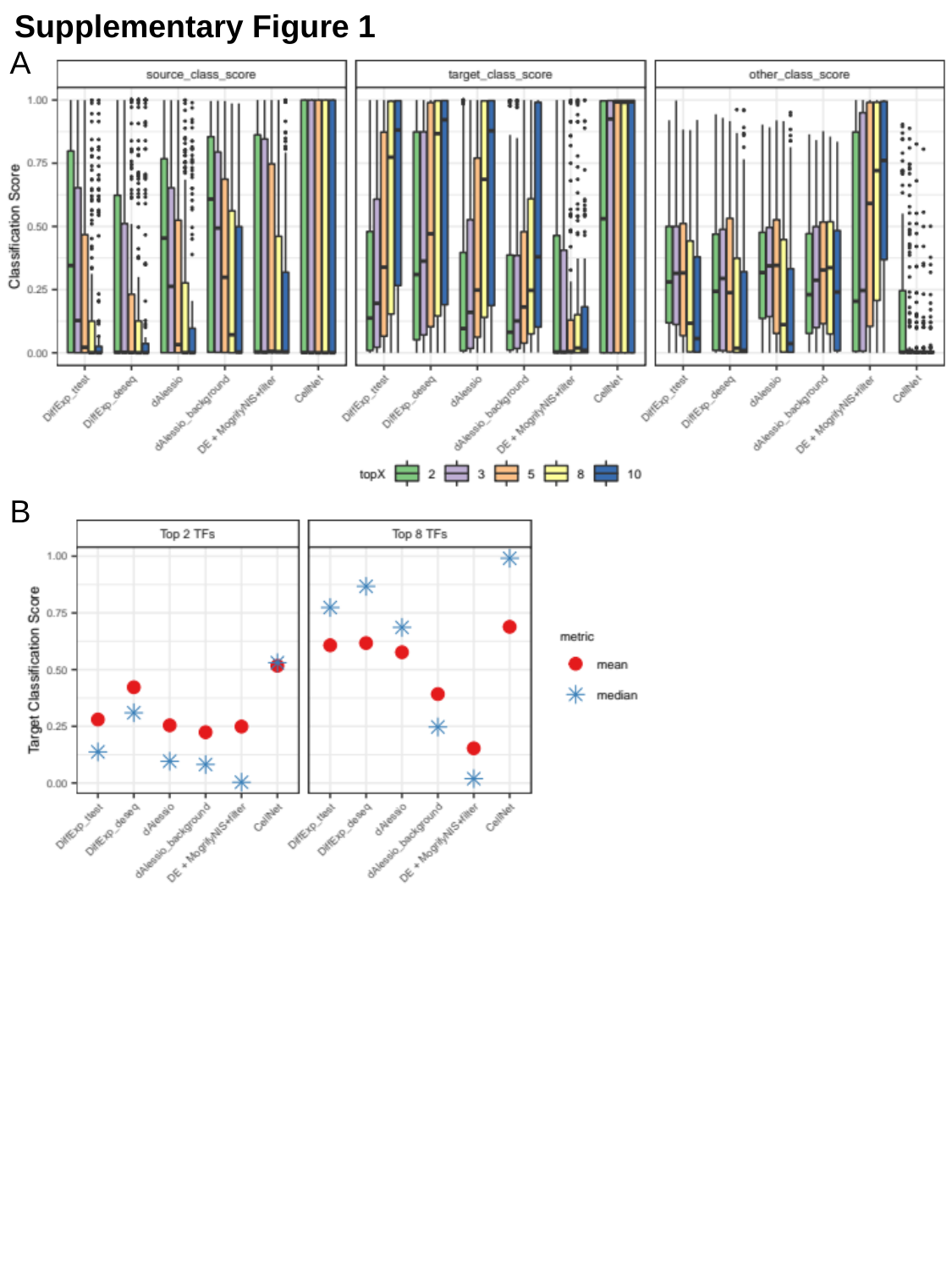

Supplementary Figure 1
A
B

### Slide 2
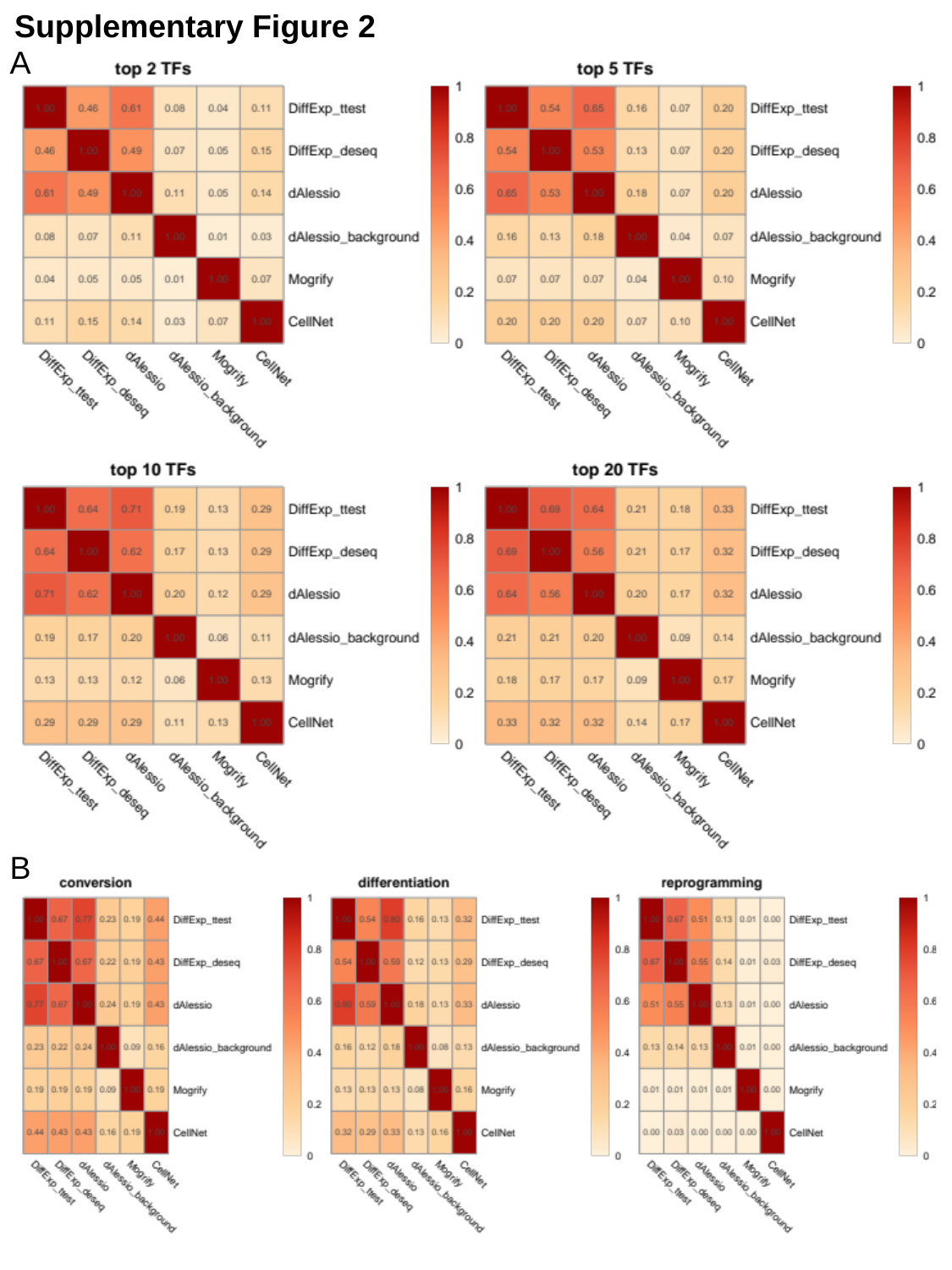

Supplementary Figure 2
A
B

### Slide 3
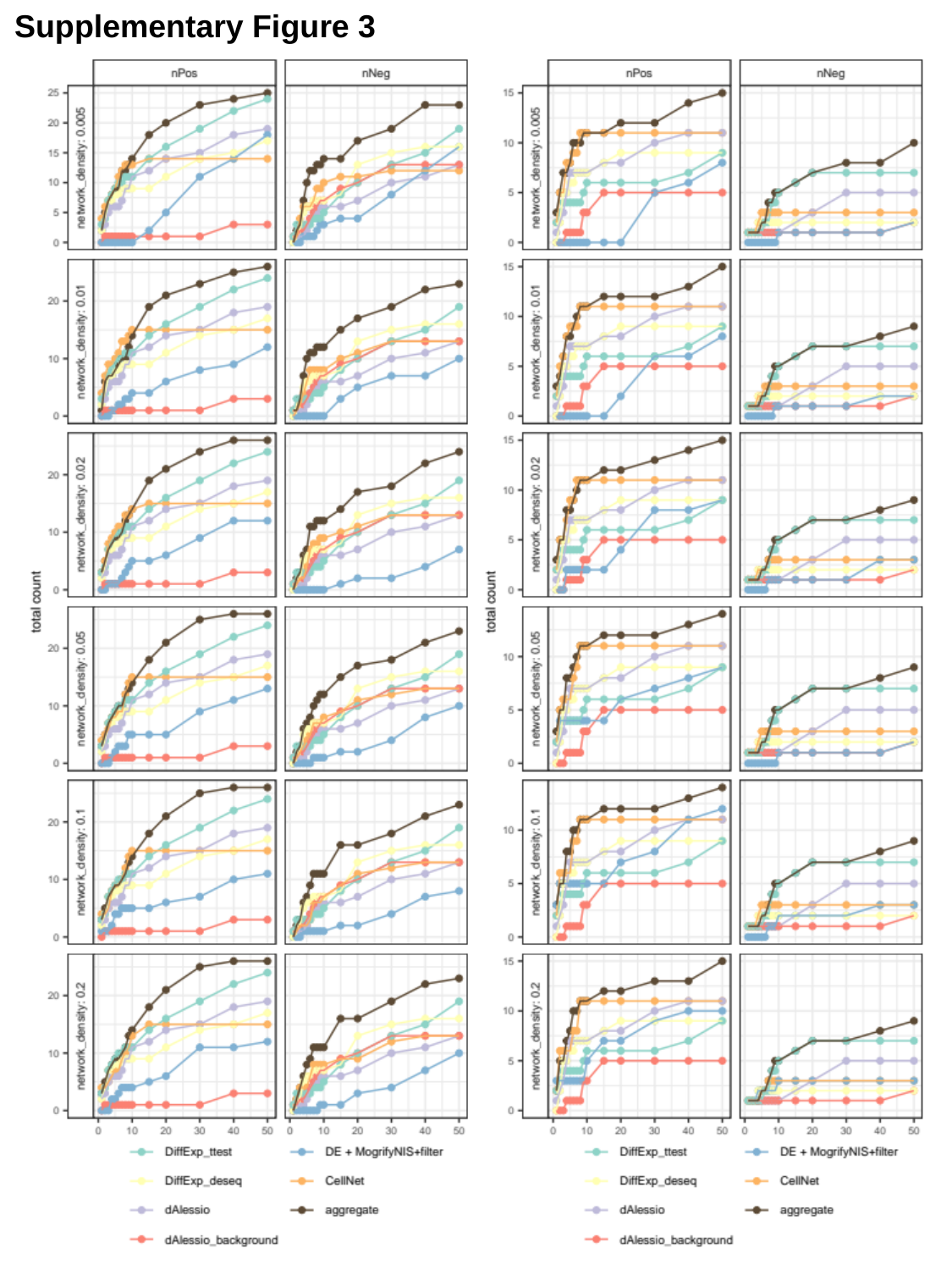

Supplementary Figure 3

### Slide 4
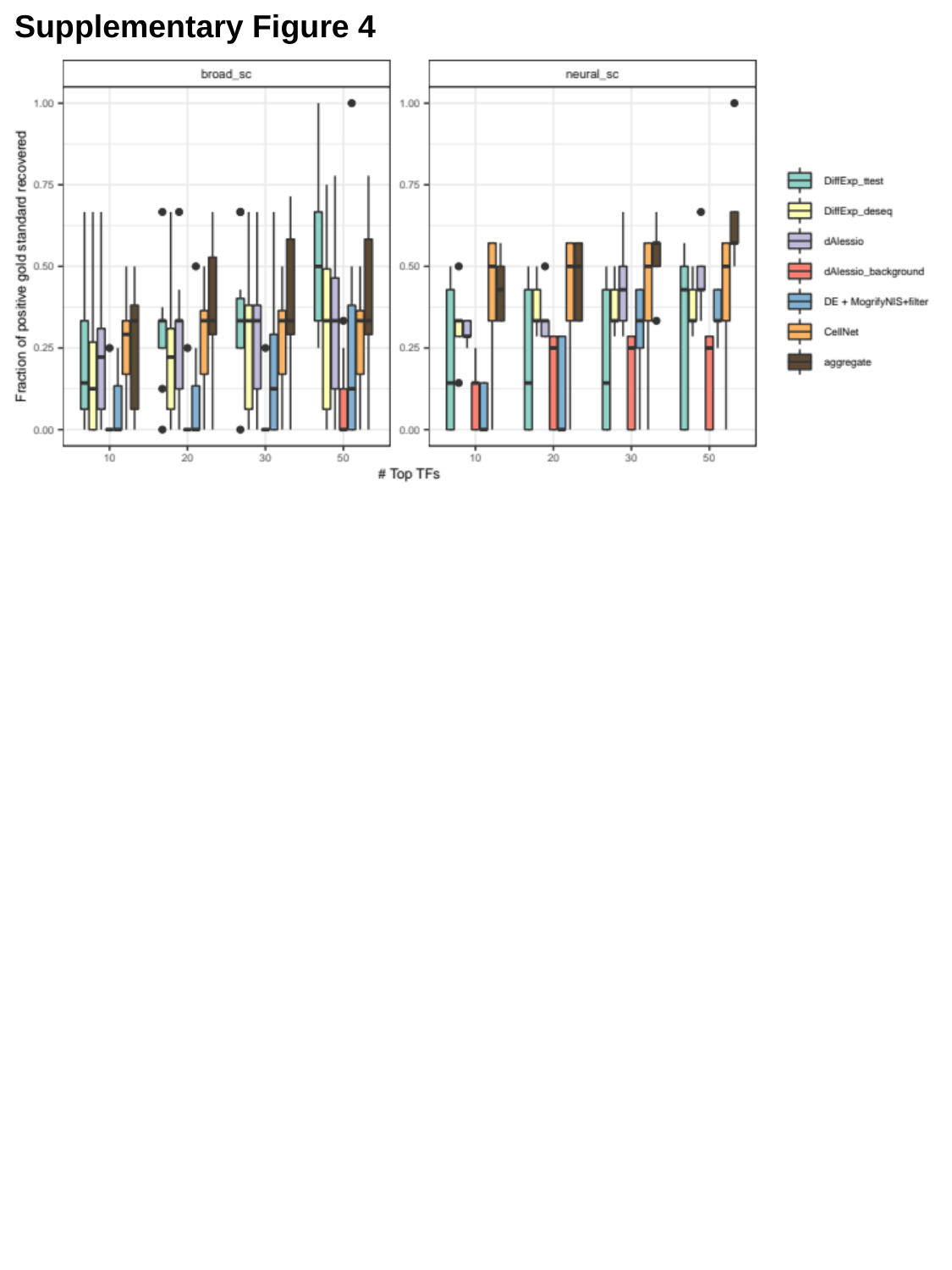

Supplementary Figure 4

### Slide 5
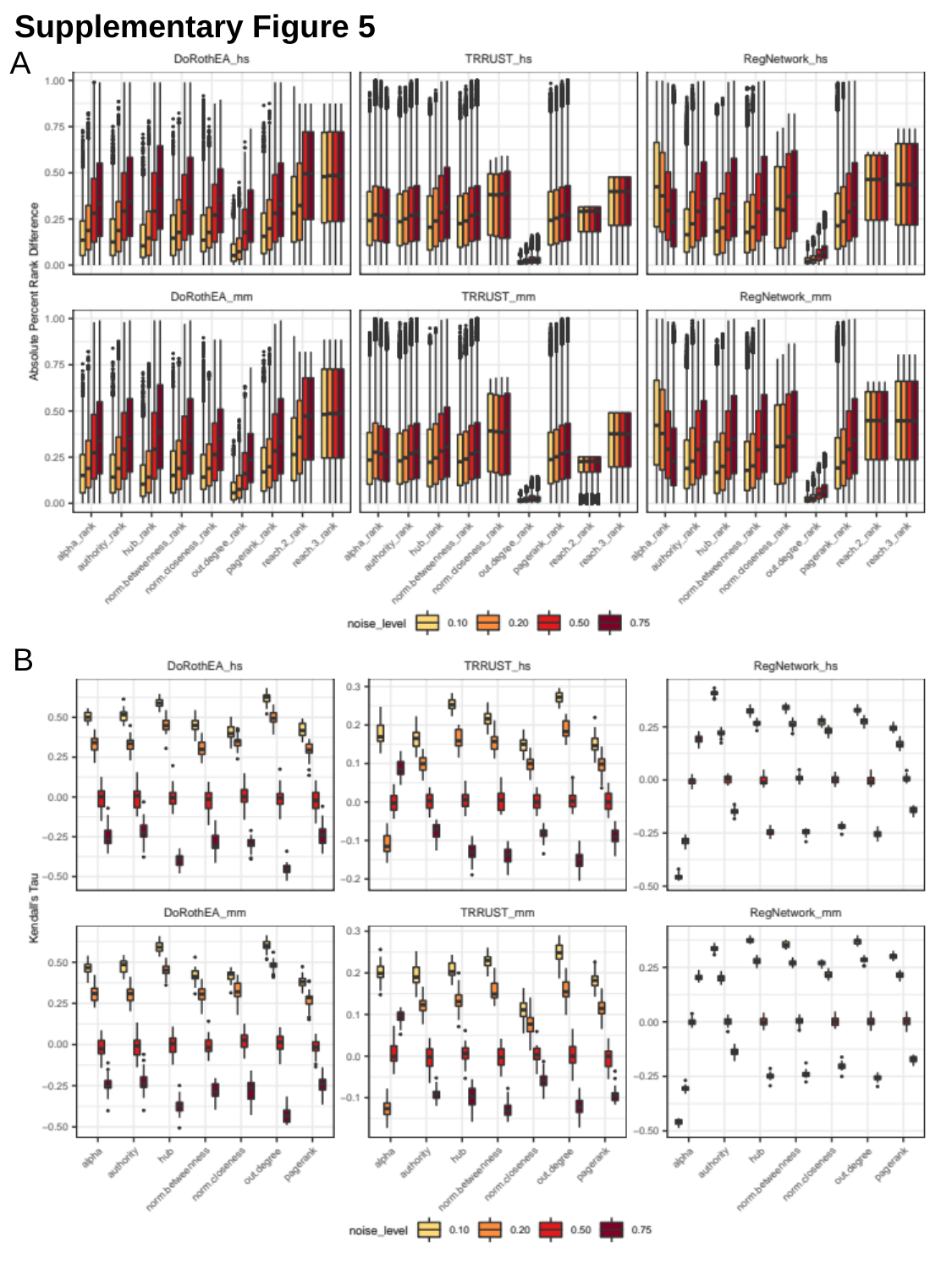

Supplementary Figure 5
A
B

### Slide 6
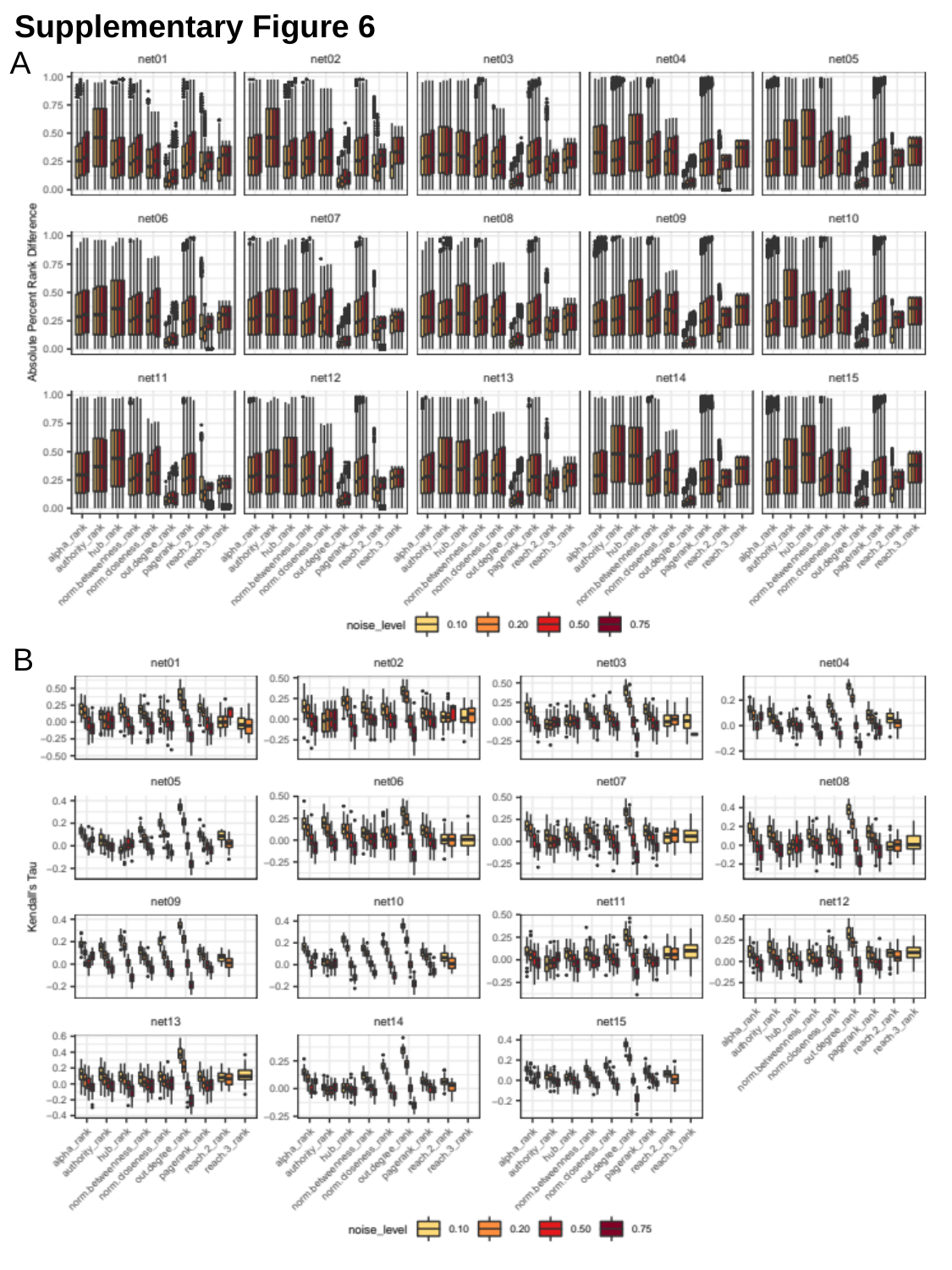

Supplementary Figure 6
A
B

### Slide 7
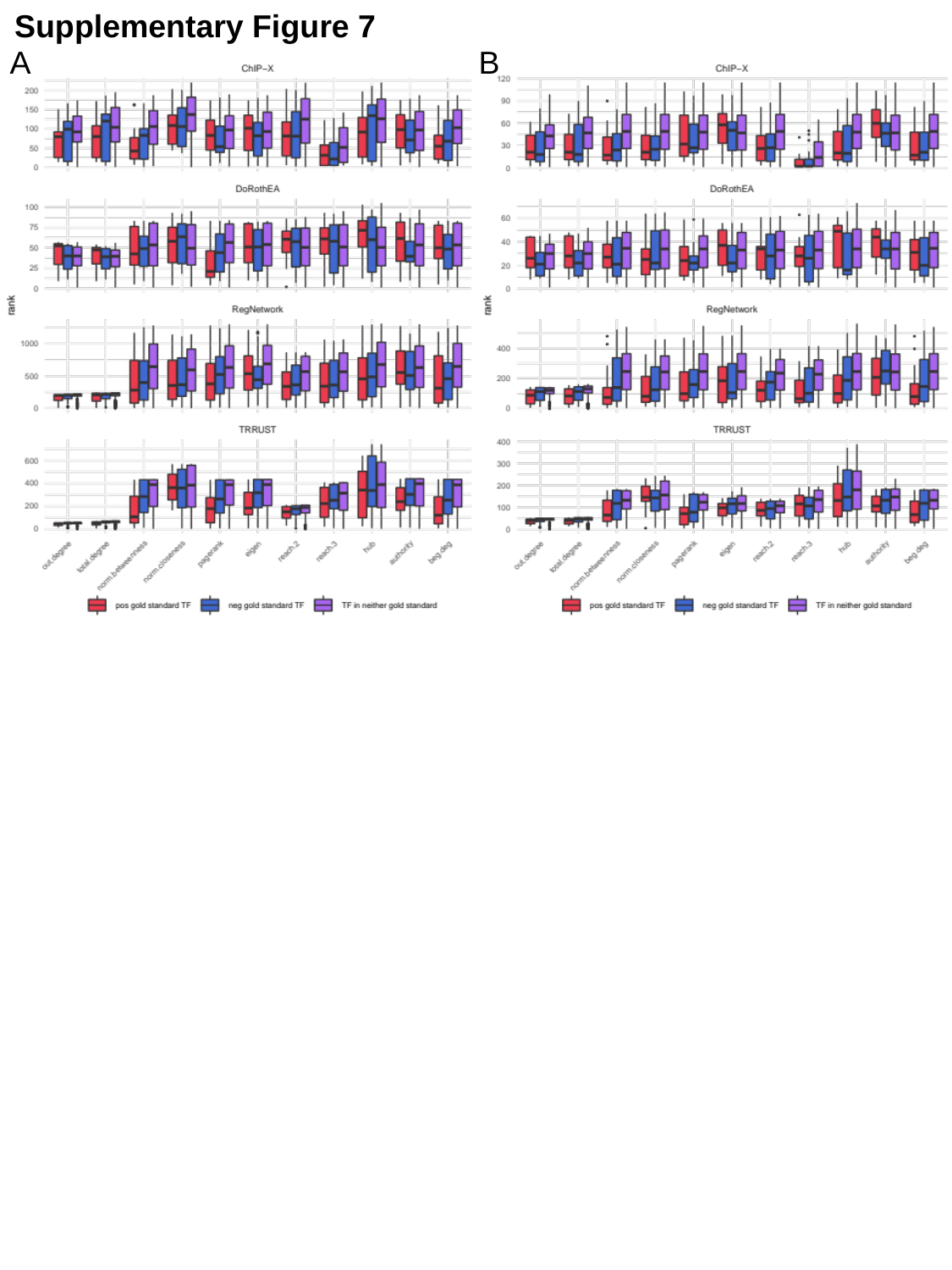

Supplementary Figure 7
A
B

### Slide 8
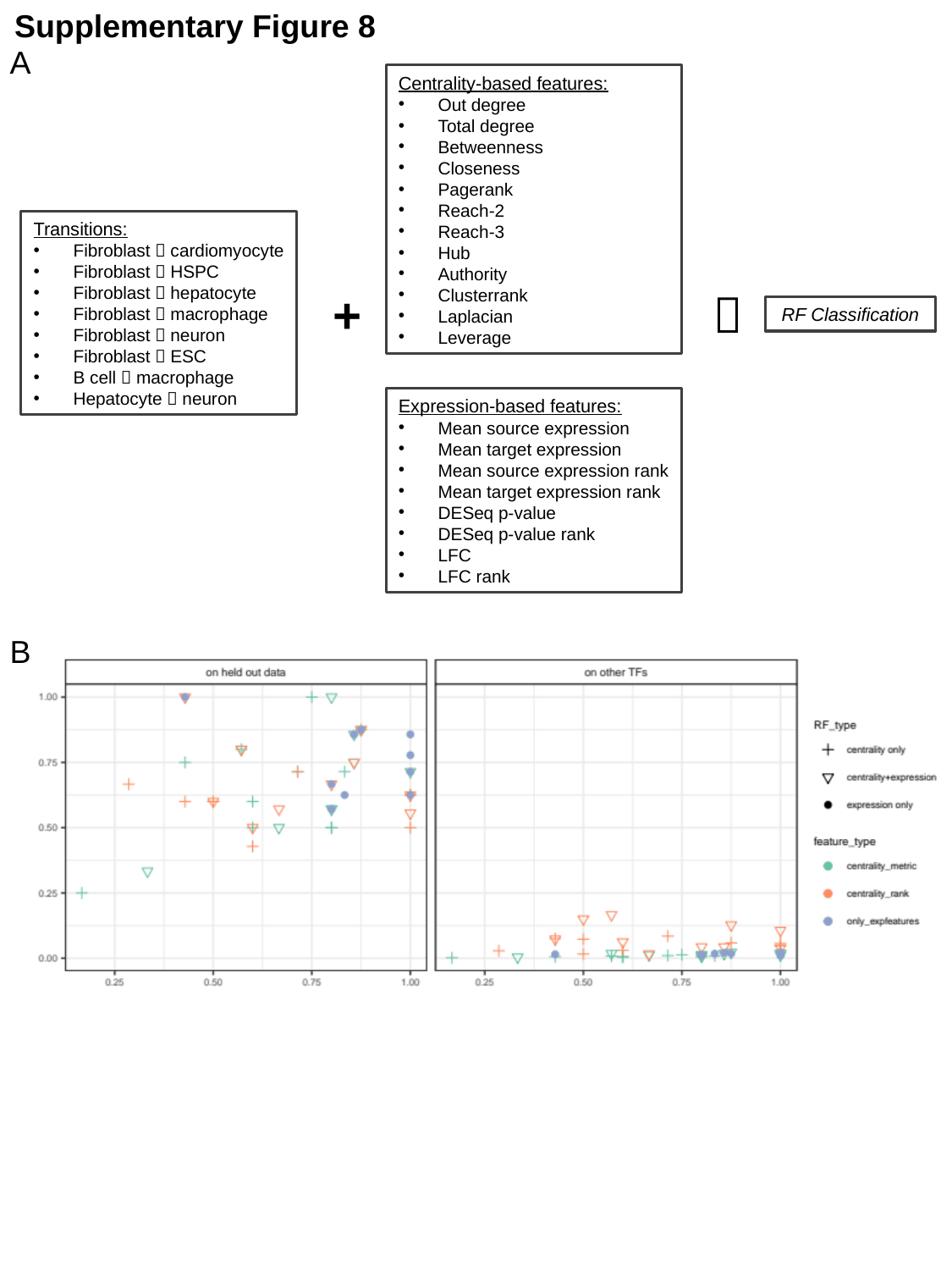

Supplementary Figure 8
A
Centrality-based features:
Out degree
Total degree
Betweenness
Closeness
Pagerank
Reach-2
Reach-3
Hub
Authority
Clusterrank
Laplacian
Leverage
Transitions:
Fibroblast 🡪 cardiomyocyte
Fibroblast 🡪 HSPC
Fibroblast 🡪 hepatocyte
Fibroblast 🡪 macrophage
Fibroblast 🡪 neuron
Fibroblast 🡪 ESC
B cell 🡪 macrophage
Hepatocyte 🡪 neuron
+
🡪
RF Classification
Expression-based features:
Mean source expression
Mean target expression
Mean source expression rank
Mean target expression rank
DESeq p-value
DESeq p-value rank
LFC
LFC rank
B

### Slide 9
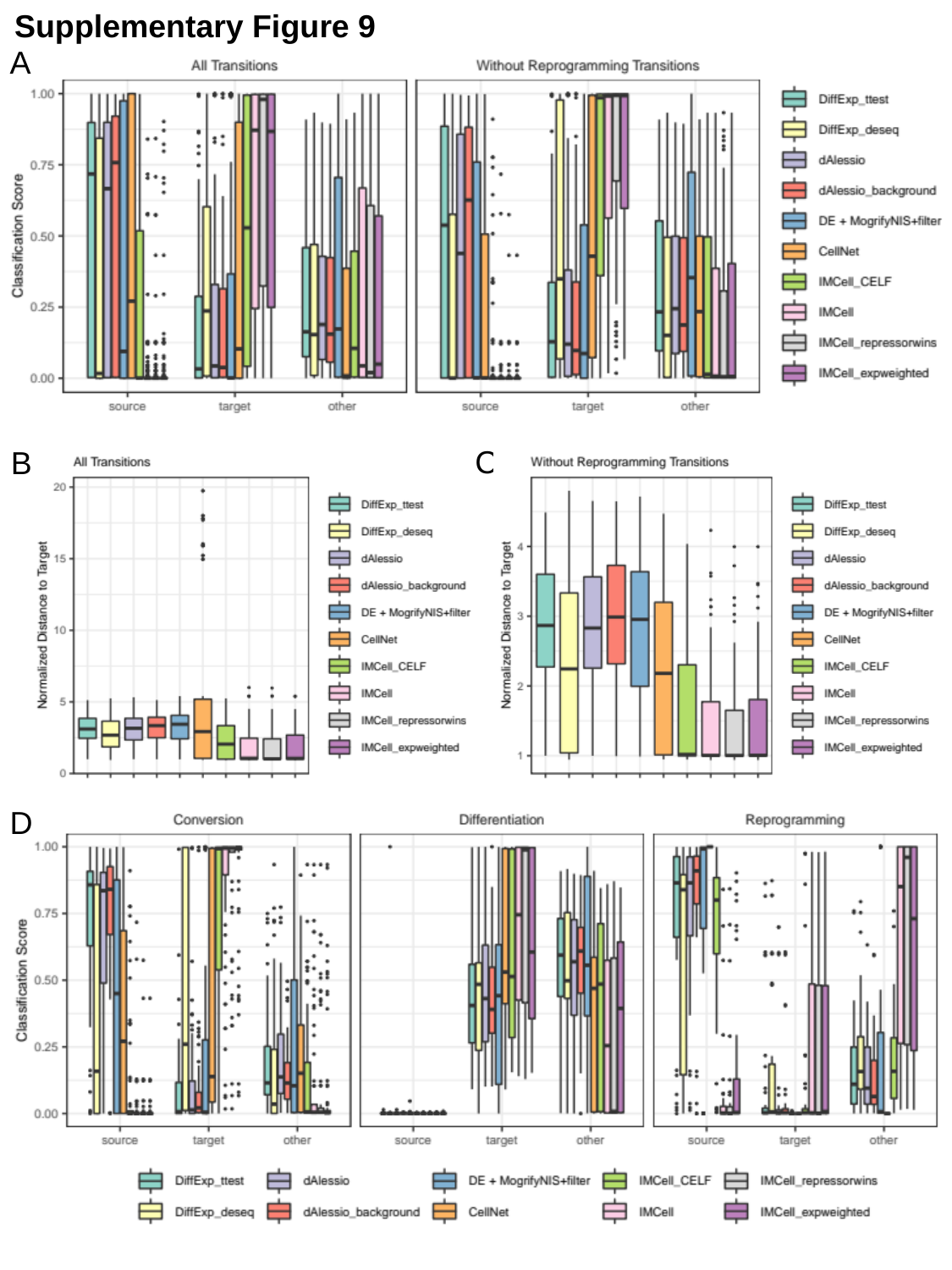

Supplementary Figure 9
A
C
B
D

### Slide 10
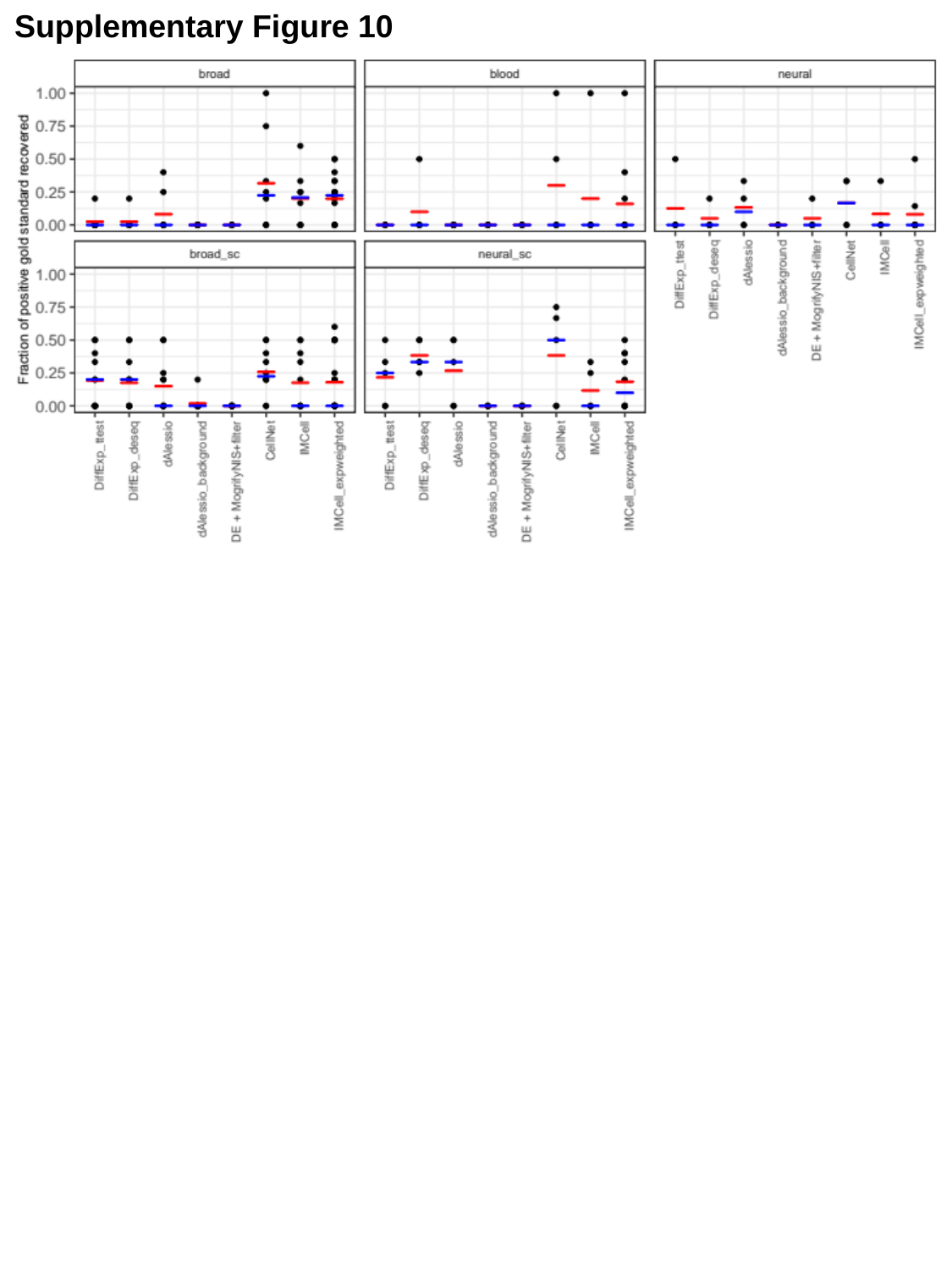

Supplementary Figure 10

### Slide 11
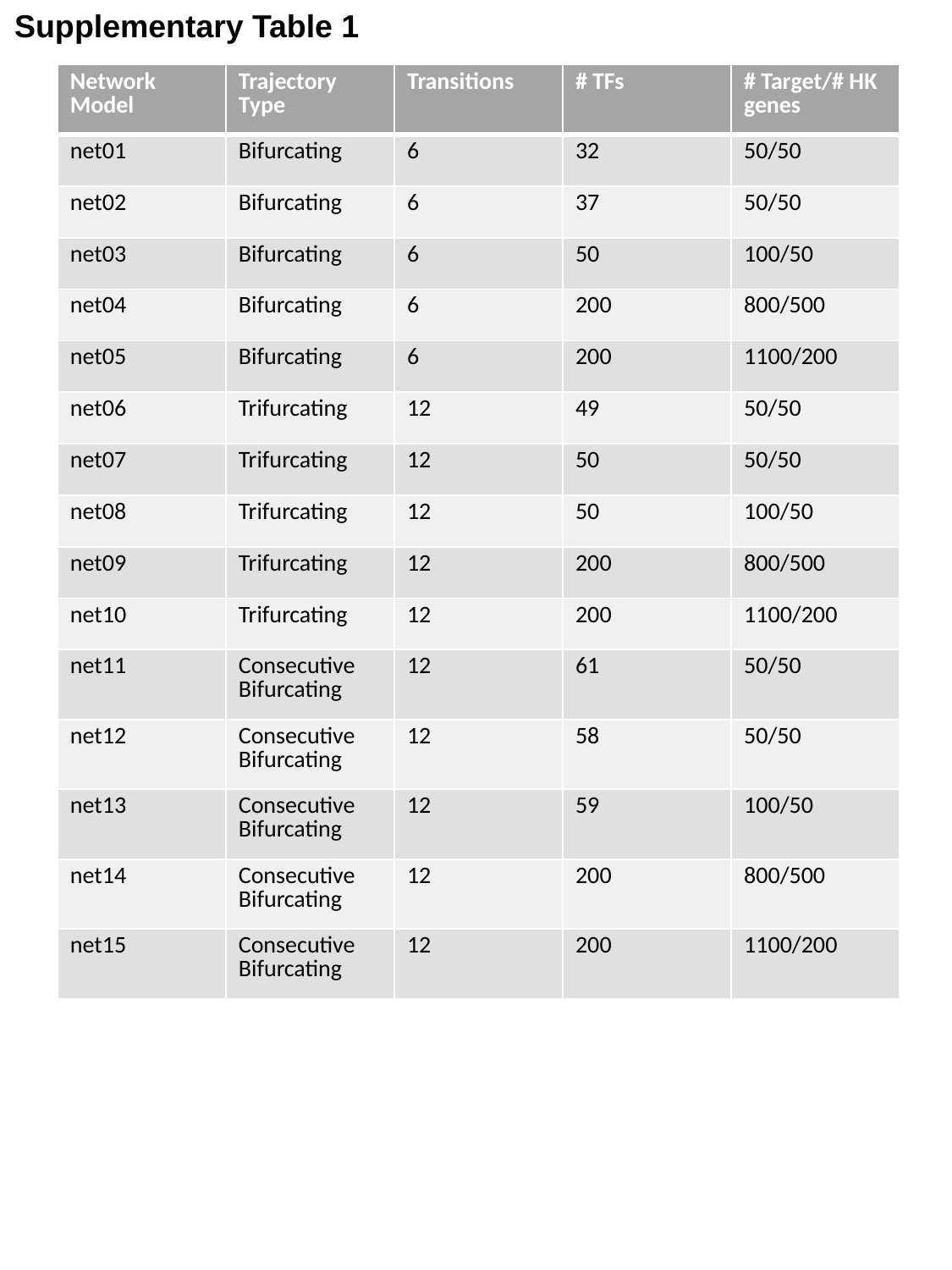

Supplementary Table 1
| Network Model | Trajectory Type | Transitions | # TFs | # Target/# HK genes |
| --- | --- | --- | --- | --- |
| net01 | Bifurcating | 6 | 32 | 50/50 |
| net02 | Bifurcating | 6 | 37 | 50/50 |
| net03 | Bifurcating | 6 | 50 | 100/50 |
| net04 | Bifurcating | 6 | 200 | 800/500 |
| net05 | Bifurcating | 6 | 200 | 1100/200 |
| net06 | Trifurcating | 12 | 49 | 50/50 |
| net07 | Trifurcating | 12 | 50 | 50/50 |
| net08 | Trifurcating | 12 | 50 | 100/50 |
| net09 | Trifurcating | 12 | 200 | 800/500 |
| net10 | Trifurcating | 12 | 200 | 1100/200 |
| net11 | Consecutive Bifurcating | 12 | 61 | 50/50 |
| net12 | Consecutive Bifurcating | 12 | 58 | 50/50 |
| net13 | Consecutive Bifurcating | 12 | 59 | 100/50 |
| net14 | Consecutive Bifurcating | 12 | 200 | 800/500 |
| net15 | Consecutive Bifurcating | 12 | 200 | 1100/200 |

### Slide 12
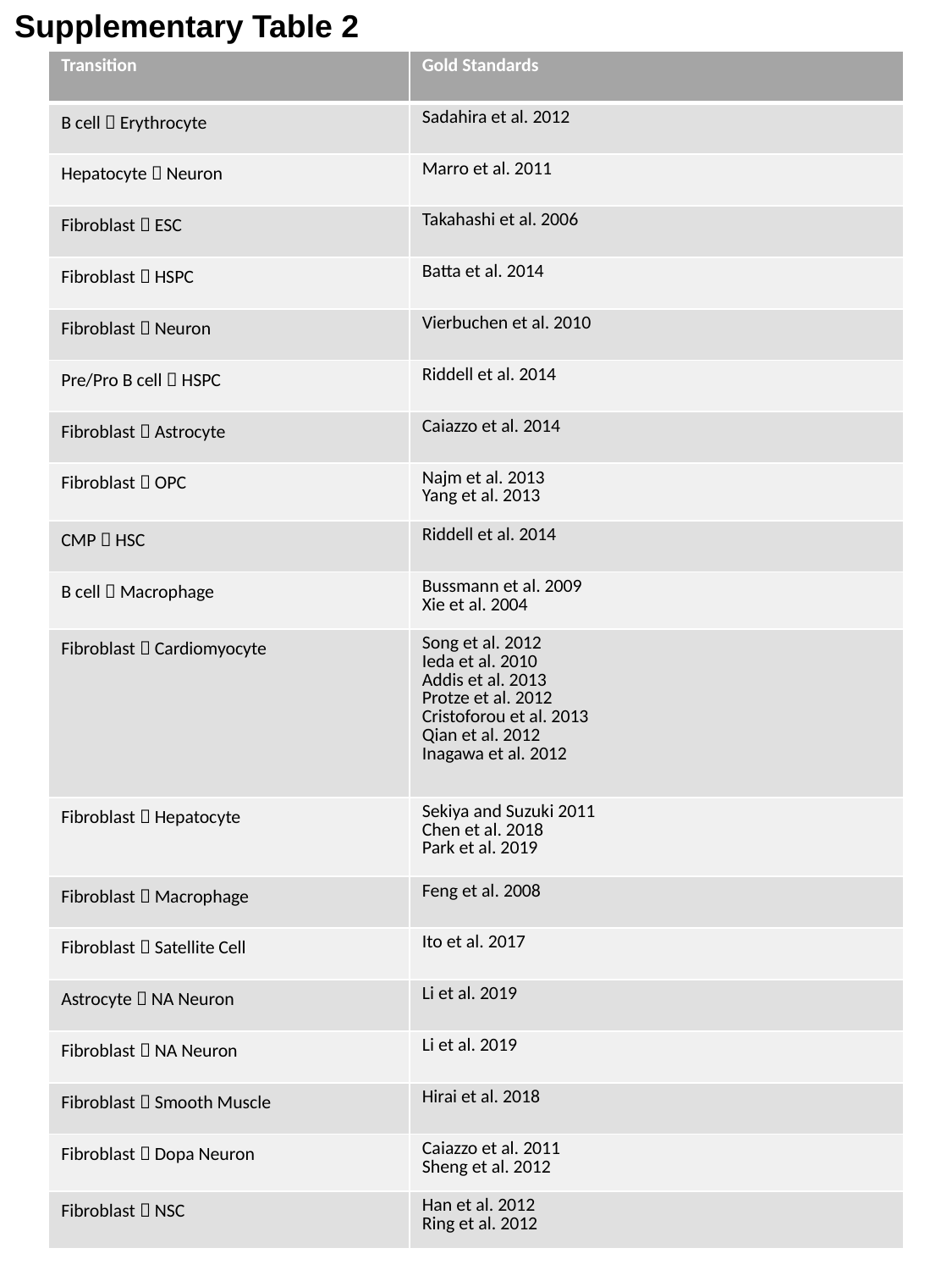

Supplementary Table 2
| Transition | Gold Standards |
| --- | --- |
| B cell 🡪 Erythrocyte | Sadahira et al. 2012 |
| Hepatocyte 🡪 Neuron | Marro et al. 2011 |
| Fibroblast 🡪 ESC | Takahashi et al. 2006 |
| Fibroblast 🡪 HSPC | Batta et al. 2014 |
| Fibroblast 🡪 Neuron | Vierbuchen et al. 2010 |
| Pre/Pro B cell 🡪 HSPC | Riddell et al. 2014 |
| Fibroblast 🡪 Astrocyte | Caiazzo et al. 2014 |
| Fibroblast 🡪 OPC | Najm et al. 2013 Yang et al. 2013 |
| CMP 🡪 HSC | Riddell et al. 2014 |
| B cell 🡪 Macrophage | Bussmann et al. 2009 Xie et al. 2004 |
| Fibroblast 🡪 Cardiomyocyte | Song et al. 2012 Ieda et al. 2010 Addis et al. 2013 Protze et al. 2012 Cristoforou et al. 2013 Qian et al. 2012 Inagawa et al. 2012 |
| Fibroblast 🡪 Hepatocyte | Sekiya and Suzuki 2011 Chen et al. 2018 Park et al. 2019 |
| Fibroblast 🡪 Macrophage | Feng et al. 2008 |
| Fibroblast 🡪 Satellite Cell | Ito et al. 2017 |
| Astrocyte 🡪 NA Neuron | Li et al. 2019 |
| Fibroblast 🡪 NA Neuron | Li et al. 2019 |
| Fibroblast 🡪 Smooth Muscle | Hirai et al. 2018 |
| Fibroblast 🡪 Dopa Neuron | Caiazzo et al. 2011 Sheng et al. 2012 |
| Fibroblast 🡪 NSC | Han et al. 2012 Ring et al. 2012 |
